# Mechanistic basis for protection against fatty liver disease by *CIDEB* loss-of-function mutations

**DOI:** 10.1101/2025.04.01.646492

**Authors:** Qiyu Zeng, Satish Patel, Xun Wang, Meng-Hsiung Hsieh, Zhijie Li, Xiongzhao Ren, Jingjing Wang, Dohun Kim, Shili Li, Xinping Gu, Greg Mannino, Gianna Maggiore, Xiangyi Fang, Lin Li, Min Zhu, Mengmeng Wang, Boyuan Li, Amaey Bellary, Koini Lim, Zhetuo Qi, Pushpa Pushpa, Mandour Omer Mandour, Vladimir Saudek, Tripti Sharma, Yu Zhang, Gerta Hoxhaj, Prashant Mishra, Purva Gopal, Peter Campbell, Matthew Hoare, David B. Savage, Hao Zhu

**Affiliations:** Children’s Research Institute, Departments of Pediatrics and Internal Medicine, Center for Regenerative Science and Medicine, Children’s Research Institute Mouse Genome Engineering Core, University of Texas Southwestern Medical Center, Dallas, TX 75390, USA; University of Cambridge Metabolic Research Laboratories, Wellcome Trust-MRC Institute of Metabolic Science, Cambridge, CB2 0QQ, UK; Department of Pathology, University of Texas Southwestern Medical Center, Dallas, TX 75390, USA; Cancer Genome Project, Wellcome Sanger Institute, Hinxton, Cambridgeshire CB10 1SA, UK; University of Cambridge Department of Medicine, Cambridge Biomedical Campus, Cambridge, CB2 0QQ, UK and University of Cambridge Early Cancer Institute, Hutchison Research Centre, Cambridge Biomedical Campus, Cambridge, CB2 0XZ, UK

## Abstract

**Background & Aims:** Somatic and germline *CIDEB* mutations are associated with protection from chronic liver diseases. The mechanistic basis and whether *CIDEB* suppression would be an effective therapy against fatty liver disease remain unclear.

**Methods:** 21 *CIDEB* somatic mutations were introduced into cells to assess functionality. In vivo screening was used to trace *Cideb* mutant clones in mice fed normal chow, western (WD), and choline-deficient, L-amino acid-defined, high-fat (CDA-HFD) diets. Constitutive and conditional *Cideb* knockout mice were generated to study *Cideb* in liver disease. Isotope tracing was used to evaluate fatty acid oxidation and de novo lipogenesis. Transcriptomics, lipidomics, and metabolic analyses were utilized to explore molecular mechanisms. Double knockout models (*Cideb/Atgl* and *Cideb/Ppara*) tested mechanisms underlying *Cideb* loss.

**Results:** Most *CIDEB* mutations showed that they impair function, and lineage-tracing showed that loss-of-function clones were positively selected with some, but not all fatty liver inducing diets. *Cideb* KO mice were protected from WD, CDA-HFD, and alcohol diets, but had the greatest impact on CDA-HFD induced liver disease. Hepatocyte-specific *Cideb* deletion could ameliorate disease after MASLD establishment, modeling the impact of therapeutic siRNAs. *Cideb* loss protected livers via increased β-oxidation, specifically through ATGL and PPARa activation.

**Conclusions:** *Cideb* deletion is more protective in some types of fatty liver disease. β-oxidation is an important component of the *Cideb* protective mechanism. *CIDEB* inhibition represents a promising approach, and somatic mutations in *CIDEB* might predict the patient populations that might benefit the most.

## Introduction

Recently, somatic genomic sequencing of chronic liver disease samples from patients with alcohol related liver disease (ALD) and metabolic dysfunction-associated steatotic liver disease (MASLD) identified *Cell Death-Inducing DFF45-like Effector Protein B* (*CIDEB*) mutations that promote clonal fitness, likely through the reduction of hepatic lipid overload ^1^. In addition, human genetic studies showed that heterozygous germline mutations in *CIDEB* protect carriers from a range of liver diseases including ALD and MASLD ^2^. Collectively, these studies suggest that a broad range of *CIDEB* mutations are protective against lipotoxic liver disease. Consistently, whole-body *CIDEB* deletion in mice reduces body weight, improves insulin sensitivity and alleviates MASLD ^3^. In keeping with the reported action of all CIDE-family proteins (CIDEA, CIDEB, CIDEC), mechanistic studies show that CIDEB directly mediates lipid droplet (LD) growth ^4^. CIDEB has also been reported to affect hepatic lipid metabolism in other ways, including de novo lipogenesis (DNL) and VLDL particle lipidation ^5–7^.

The recent human genetic data have generated considerable interest in *CIDEB* as a therapeutic target for MASLD because of its growing prevalence and association with end-stage liver disease ^8^. Although whole-body *Cideb* deletion in mice appears to be beneficial, it is unclear if and how *CIDEB* somatic mutant clones are selected for within fatty livers, and if liver-specific loss of *CIDEB* would also protect against established MASLD. Furthermore, the molecular basis for CIDEB’s effects on liver biology remain incompletely understood ^9^.

Here, we sought to understand the molecular basis for the selection of human *CIDEB* somatic mutations by exploring structure/function relationships. In addition to demonstrating that these mutants impair LD growth to at least some extent, these studies suggest that CIDEB oligomerization can result in the formation of a hydrophobic channel between adjacent LDs. We then asked if *Cideb* deficient clone selection was more pronounced in certain types of liver disease, and determined if this information might provide clues about the molecular basis for CIDEB’s effects. We asked if the degree of positive clone selection is correlated with the therapeutic efficacy of liver-wide *Cideb* deletion. These studies showed that CIDEB predominantly facilitates unsaturated fatty acid accumulation in LDs, whereas its deficiency enhances fatty acid oxidation (FAO) in the liver. Finally, therapeutic modeling shows that liver-specific *CIDEB* suppression can ameliorate late stage manifestations of the disease such as fibrosis and liver cancer. These phenotypic and mechanistic findings pave the way for biomarker-based therapeutic studies using liver-specific targeting of *CIDEB*.

## Results

### Functional characterisation of somatic *CIDEB* mutations

As all CIDE-family proteins facilitate LD growth via formation of contact sites between the surface phospholipid monolayers of adjacent LDs ^4,9–12^, we sought to measure the effects of *CIDEB* mutants on LD size *in vitro*. *CIDEB* missense mutations detected by somatic sequencing spanned all structurally validated and AlphaFold predicted domains (**Fig. S1**) ^1^. Among the mutations, two (W28* and W181*) were predicted to result in premature stop codons and likely nonsense mediated decay, whereas one distal frameshift mutant (Y220fs) was not predicted to undergo nonsense mediated RNA decay. When expressed in oleate loaded HeLa cells, expression of W28* and Y220fs were severely reduced whereas W181* was highly expressed (**Fig. 1A**). However, all three clearly inhibited LD growth (**Fig. 1B,C**). All 18 tested missense variants were expressed at the protein level in HeLa cells, though some mutations, such as the E78K mutant, did reduce protein expression (**Fig. 1A**). While there was substantial variation in the impact on LD size, none significantly enhanced LD enlargement and 10 out of 18 missense variants significantly reduced LD size compared with wild-type (WT) *CIDEB* (**Fig. 1B**). LD targeting for most variants remained unperturbed, but 3 that did not reduce LD size (L4H, L7Q, K62E) showed more cytosolic distribution and reduced protein expression (**Fig. 1A,C**).

**Fig. 1.**
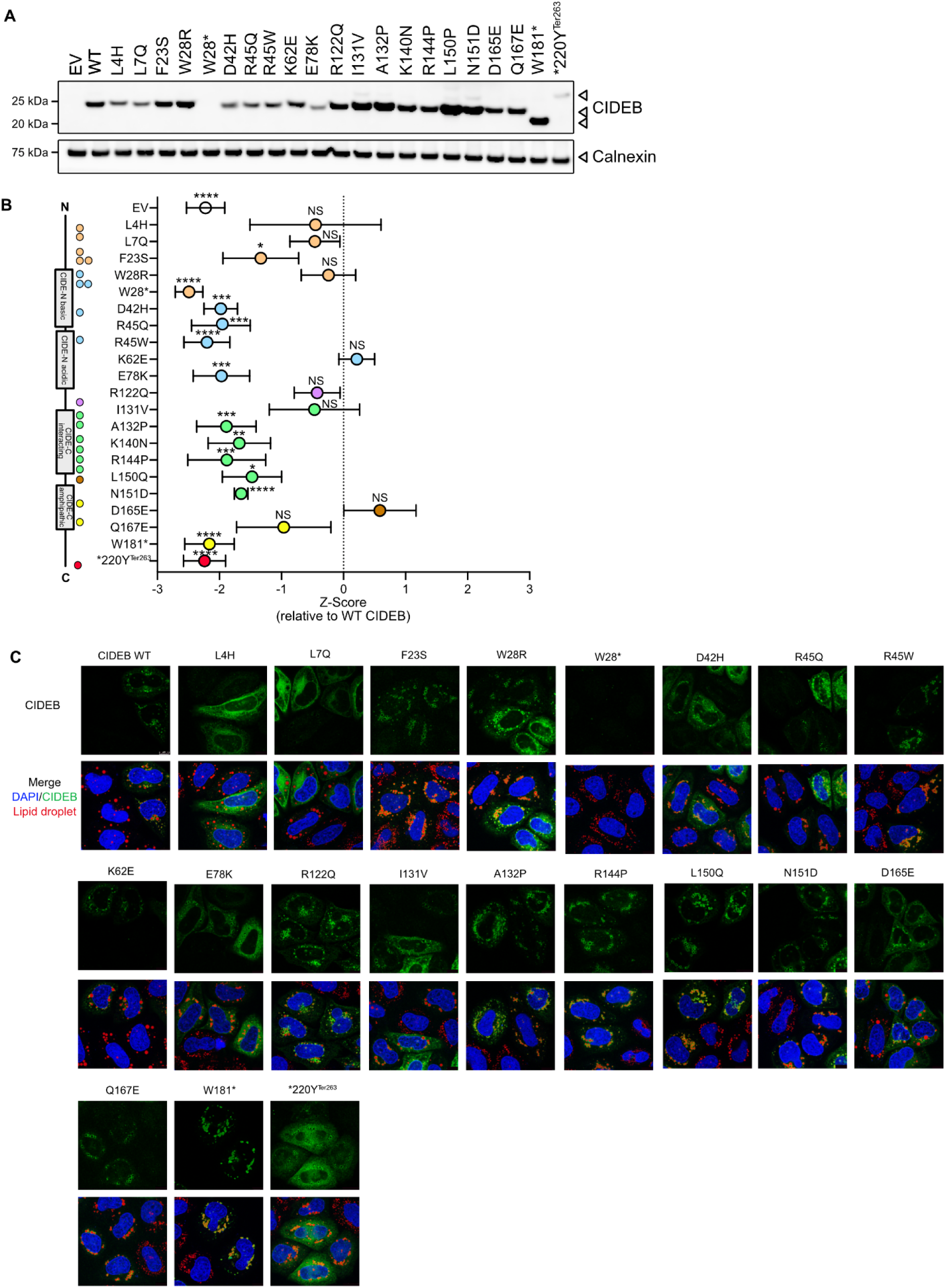
Impact of somatic *CIDEB* mutations on expression and LD enlargement. **A.** Western blot analysis in oleate-loaded HeLa cells transfected with *CIDEB*. This blot is representative of 3 independent experiments. **B.** Quantification of LD volume in oleate-loaded HeLa cells transfected with *CIDEB* mutants. Enlargement activity is represented as a Z-score where 0 is WT *CIDEB* **C.** Representative images of LDs (red) and CIDEB (green) in oleate-loaded HeLa cells transfected with *CIDEB*. Nuclei were stained with DAPI (blue).

The structure of the globular CIDE-N domain (located at the amino terminus of the CIDE family of proteins) was previously resolved ^13^ and recent studies have characterized the amphipathic helical domain ^11^. Because the predicted β-sheet region of the CIDE-C domain (located at the carboxy terminus) is less well studied, we next focused on this domain. Interestingly, AlphaFold predicted that it can fold into a tetrameric β-barrel with a central hydrophobic tunnel (**Fig. 2A**). This would effectively lead to formation of a channel (**Fig. 2B)** with a hydrophilic outer surface (**Fig. 2C)** and a hydrophobic pore (**Fig. 2D**), potentially enabling lipid transfer between LDs. To test this model, we designed two ‘artificial’ missense mutants (F134D and V152D) to replace hydrophobic residues inside the putative channel with charged amino acids (**Fig. 2E**). Consistent with the predicted model, both mutations significantly impaired LD growth (**Fig. 2F-H)**. Parallel mutagenesis experiments with CIDEC showed similar results (**Fig. S2A-C**). The proposed channel is hydrophobic on the inside, which would allow the transfer of neutral lipids.

**Fig. 2.**
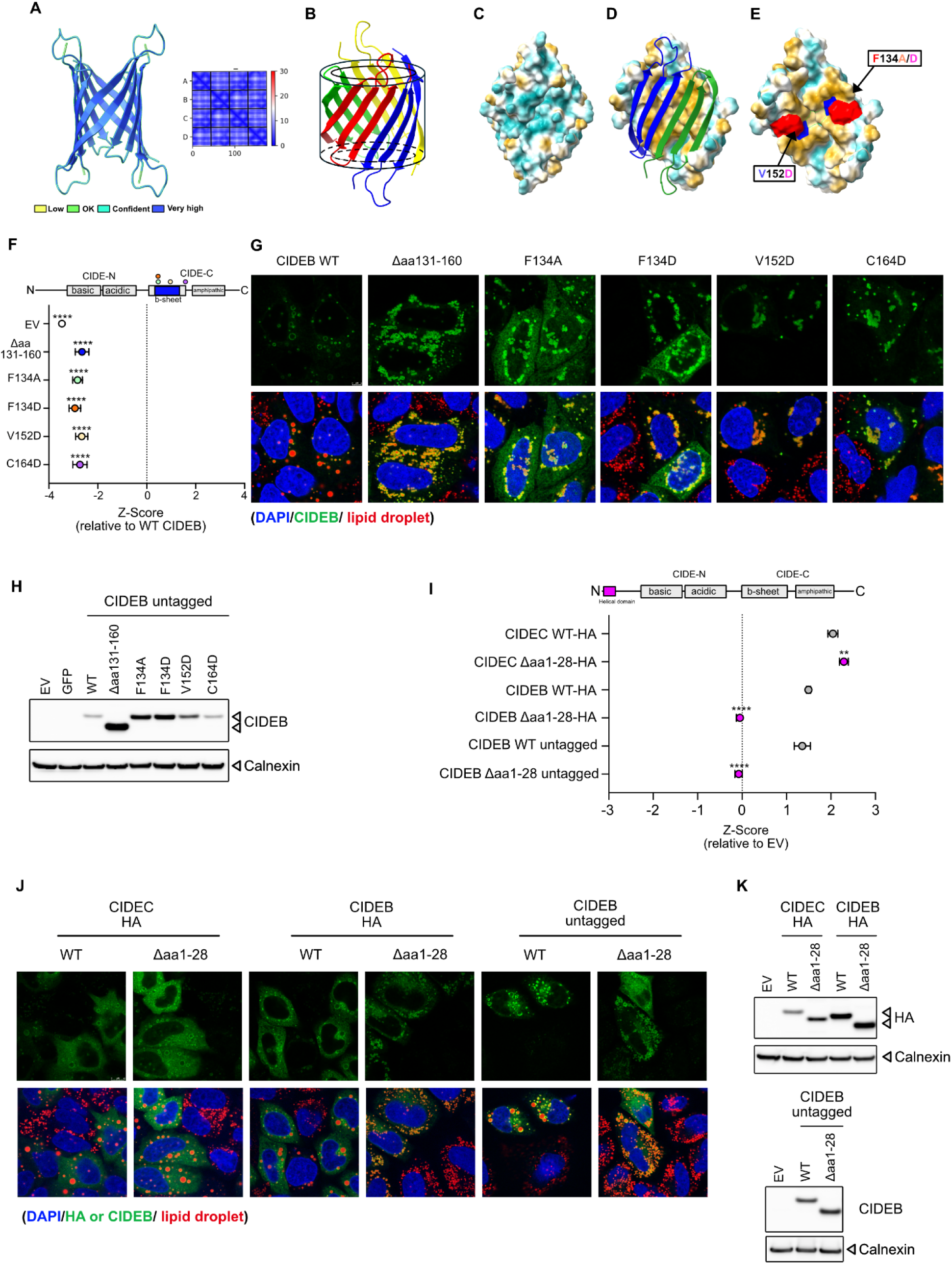
Disruption of the C-terminal β-sheet and N-terminus helix affects CIDEB-mediated LD enlargement. **A.** AlphaFold prediction of the CIDEB β-sheet aa 127-170 tetramer (left panel). Predicted structure coloured according to predicted local distance difference (Quality parameters displayed below). Predicted aligned error (Å) between residues of the four protomers A to D, coloured according to scale on the left (right panel). Black lines separate the sequences of the monomeric units. **B.** Model of CIDEB aa 127-170 forming a hollow β-barrel. The four protomers are rendered in different colors. **C.** Surface representation of the external hydrophilic surface of the CIDEB putative channel. **D.** Surface representation of the inner hydrophobic surface of the CIDEB putative channel. The two protomers are presented as ribbons to render the interior visible. **E.** Surface representation of CIDEB putative channel. F145 (red) and V152 (blue) at the inner surface (two protomers shown). Lipid transfer is suppressed when mutated to D. The residues between the two V152 from two neighbouring protomers are C164 (green contour, **Fig. S2**). The Intensity of yellow and cyan corresponds to the degree of hydrophobicity and hydrophilicity, respectively. **F.** Quantification of LD volume in oleate-loaded HeLa cells transfected with CIDEB β-sheet deletion and point mutants.Enlargement activity is represented as a Z-score where 0 is WT *CIDEB* **G.** Representative images of LDs (red) and CIDEB (green) in oleate-loaded HeLa cells transfected with CIDEB β-sheet deletion and point mutants. Nuclei were stained with DAPI (blue). **H.** Western blot analysis in oleate-loaded HeLa cells transfected with CIDEB β-sheet deletion and point mutants. **I.** Quantification of LD volume in oleate-loaded HeLa cells transfected with wildtype (WT) or n-terminal helix deleted mutants (aa1-29) of HA-tagged CIDEC, HA-CIDEB or untagged CIDEB. LD enlargement activity is represented as a Z-score where 0 is empty vector (EV). **J.** Representative images of LDs (red) and HA (green) or CIDEB (green) in oleate-loaded HeLa cells transfected with wildtype (WT) or n-terminal helix deleted mutants (aa1-29) of HA-tagged CIDEC, HA-CIDEB or untagged CIDEB. Nuclei were stained with DAPI (blue). **K.** Western blot analysis in oleate-loaded HeLa cells transfected with HA tagged CIDEC, HA tagged CIDEB, or untagged CIDEB n-terminal helix deleted mutants. All data are presented as mean ± SEM ( **p < 0.01; ****p < 0.0001).

Analysis of the evolution of all CIDE proteins showed that none of them have charged residues inside this putative channel (**Fig. S2D**). We further evaluated the stability of the inter-protomer β-barrel, which appears to be stabilized by π-π aromatic interactions inside the channel (**Fig. S2E**). In keeping with this observation, a missense F134A mutant that interrupts the aromatic interaction reduced LD growth (**Fig. 2F-H**). In addition, the model places C164 in two antiparallel protomers in an ideal position to form a stabilizing disulfide bond (**Fig. 2E** and **Fig. S2F**), and introducing a mutation C164D prevented CIDEB from enlarging LDs (**Fig. 2F-H**).

Four (K140N, R144P, L150Q, N151D) of the human somatic CIDEB variants in the β-sheet region substantially impaired LD growth whereas two (D165E and Q167E) did not (**Fig. 1B**). K140 and L150 are conserved in all CIDE proteins, whereas R144 is only conserved in CIDEA and CIDEB. Our tetrameric β-barrel model places K140 and R144 in the loop connecting the first and second β-strand of the putative channel, two at each entry. They could interact with the negative phospholipid surface of LDs. The mutations remove the positive charge and thus could abolish the interaction. L150 is a hydrophobic residue inside the channel. The mutation L150Q introduces a polar residue that could hinder the passage of lipids similarly to our designed mutations F134D and V152D. The tetrameric β-barrel model does not suggest an explanation for N151D introducing a positive charge at the external surface of the channel. N151 could be a glycosylation site and glycosylation is known to be important for CIDE protein function ^14^, although the specific sites have not yet been reported. The neutral mutations D165E (conserved) and Q167E (variable) at the external surface are predicted to bring about very mild changes.

Another region highlighted by the human genetic data was the extreme amino-terminus of CIDEB. In CIDEC, this region (amino acids 2-33) is predicted to fold into a short helix with no known function (**Fig. S2G**). However, deleting it entirely failed to perturb CIDEC action on LD growth. In CIDEB, both the somatic missense variants affecting this ‘domain’ (L4H and L7Q) very modestly reduced LD growth (**Fig. 1B**) but, in contrast to CIDEC, deletion of this region severely reduced CIDEB mediated LD growth (**Fig. 2I-K**). The predicted structure of this helix in CIDEB displays a continuous hydrophobic ridge on one side (L12, L13, V16, I19, F23, V27) with all charged residues on the other side (R14, R25, R26, E22) (**Fig. S2H**). The F23S mutation interrupted this bridge and significantly reduced LD size (**Fig. 1B**). The hydrophobicity of the residues in the ridge, though not the specific amino acid identity, is conserved throughout evolution (**Fig. S2**). The charged residues are conserved, with R14 replaced by K in some species. The hydrophobic ridge can also be seen in CIDEA and CIDEC. While E22 and R26 conservation is unique to CIDEB, R14 in CIDEC and R25 in CIDEA are also conserved. The pattern of conservation suggests a functional role for this helix, which could interact with other proteins. Collectively, these data suggest that the somatic CIDEB mutants are loss-of-function and that there is considerable variation in the extent of functional impairment. Specific mutations also deepen understanding of the fundamental mechanisms by which CIDE-family proteins facilitate LD growth. We proceeded to further characterize the impact of *Cideb* deletion *in vivo*.

### *Cideb* loss-of-function clones are positively selected in some fatty liver models

In order to understand the conditions under which *CIDEB* loss-of-function clones are positively selected for in liver tissues, we used two diets known to induce different forms of MASLD: a western diet (WD), and a choline-deficient, L-amino acid-defined, high-fat diet (CDA-HFD) ^15^. Compared to mice fed with a WD for 2 weeks or 12 weeks, those fed with a CDA-HFD manifested higher liver/body weight ratios (**Fig. 3A,B** and **Fig. S3A**) and increased steatosis (**Fig. 3C** and **Fig. S3B**) despite lower body weights. *Cideb* mRNA levels in the liver remained relatively stable across the various diets (**Fig. S3C,D**), while *Cidea* and *Cidec* increased in both MASLD diets (**Fig. S3E**).

**Fig. 3.**
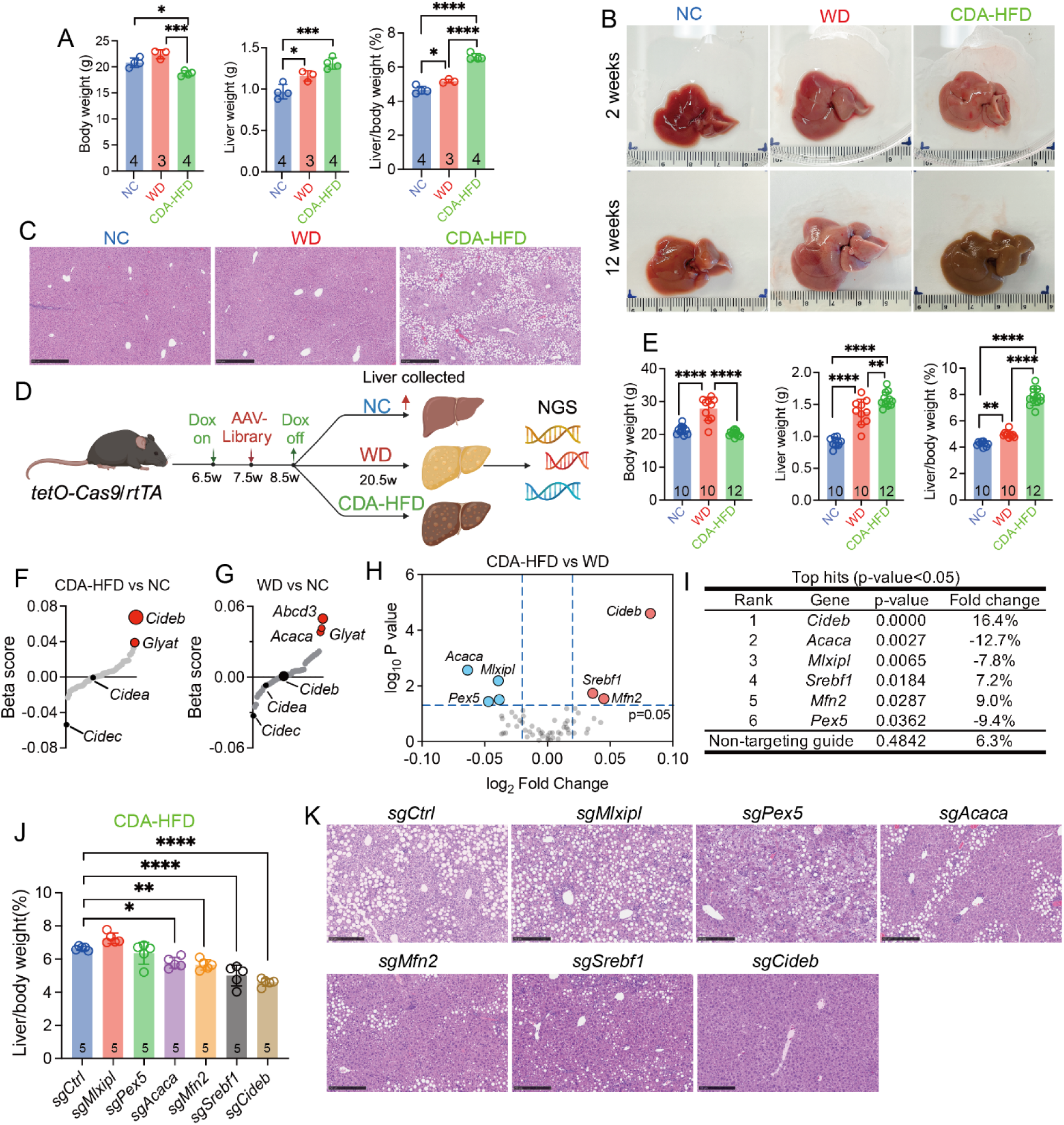
Positive selection of *Cideb* loss-of-function clones in two fatty liver models. **A.** Body weight, liver weight, and liver/body weight ratio in WT mice fed with NC, WD, or CDA-HFD for 2 weeks. **B.** Representative gross livers treated for 2 and 12 weeks on various diets. **C.** Representative H&E staining of livers from Fig. 3A. **D.** Schematic of the MOSAICS system. Cas9 was induced in iCas9 male mice using dox (1 mg/mL) from 6.5-8.5 weeks of age. AAV-sgRNA libraries targeting 55 genes were injected at 7.5 weeks of age. Mice were then fed with NC, WD, or CDA-HFD for 3 months (n = 10, 10, and 12 mice). Liver samples were collected for genomic DNA extraction and amplicon sequencing. **E.** Body weight, liver weight, and liver/body weight ratio of screening mice from Fig. 3D. **F.** β-scores from mice on CDA-HFD versus NC. Positive β-scores indicate sgRNA enrichment in the CDA-HFD group, while negative scores indicate sgRNA non-enrichment relative to NC. **G.** β-scores from mice on WD versus NC. **H.** Volcano plot comparing WD and CDA-HFD treatments, revealing enrichment of sgRNAs against *Acaca*, *Mlxipl*, and *Pex5* in WD, and *Cideb*, *Srebf1*, and *Mfn2* in CDA-HFD. **I.** Genes associated with sgRNAs in **h** with significant enrichment (p < 0.05). **J.** Validation studies shown here. Cas9 was induced from 6.5-8.5 weeks of age. At 7.5 weeks, mice received a high dose of AAV-sgRNAs (*sgCtrl*, *sgMlxipl*, *sgPex5*, *sgAcaca*, *sgMfn2*, *sgSrebf1*, or *sgCideb*) to delete genes across the liver. At 8.5 weeks, mice were fed CDA-HFD for 3 weeks. Liver/body weight ratio is shown here. **K.** Representative H&E staining of livers from **j**. All data are presented as mean ± SEM (*p < 0.05; **p < 0.01; ***p < 0.001; ****p < 0.0001).

To compare the impact of *Cideb* deficient clones with that of other genes/proteins involved in lipid metabolism, we employed the Method Of Somatic AAV-transposon *In vivo* Clonal Screening (MOSAICS) that was previously engineered to generate and track the fates of somatic mutations within the liver (**Fig. 3D**). MOSAICS can monitor the selection of mutant clones under different environmental conditions ^16^. To compare *Cideb* with other genes, we designed an AAV-sgRNA library targeting 55 genes including LD, DNL, FAO, and peroxisome genes (genes listed in **Fig. S4A**). After creating mosaic livers by injecting an AAV-sgRNA library into dox-inducible Cas9 mice, we fed the mice with normal chow (NC), CDA-HFD, or WD for 12 weeks before sequencing liver samples and comparing the sgRNA abundance between the AAV library and the harvested livers (**Fig. 3D**). The liver and body weights were similar to those from mice that were not given AAV-sgRNA libraries (**Fig. 3E** and **Fig. S3A**).

Comparing livers treated with CDA-HFD and NC revealed that clones with sgRNAs targeting *Cideb* and *Glyat* were most enriched in CDA-HFD livers (**Fig. 3F**). *Glyat*, or glycine N-acyltransferase, conjugates glycine with various acyl-CoA substrates to facilitate their excretion ^17^. In contrast, comparing livers treated with WD and NC revealed that sgRNAs against *Acaca*, *Abcd3*, and *Glyat* were enriched whilst sgRNAs against *Cidea, Cideb, or Cidec* were not positively selected (**Fig. 3G**). ACACA (ACC1) converts acetyl-CoA to malonyl-CoA during fatty acid synthesis and ABCD3 transports molecules across peroxisomal membranes ^18^.

Comparing the screens using CDA-HFD with WD revealed that sgRNAs against *Cideb, Srebf1,* and *Mfn2* were specifically enriched in CDA-HFD but not WD livers (**Fig. 3H,I**). SREBP1c is the master transcriptional regulator of lipogenesis ^19^. MFN2, plays a role in mitochondrial fusion and regulates ER stress in the liver ^20^. Conversely, sgRNAs against genes such as *Acaca, Mlxipl* and *Pex5* were significantly enriched in WD but not in CDA-HFD (**Fig. 3H,I** and **Fig. S4B**). MLXIPL, or ChREBP, is a transcription factor that regulates fatty acid synthesis in response to changes in carbohydrate metabolism ^21^. PEX5 is a receptor for peroxisomal targeting sequences involved in protein import into peroxisomes ^22^. To determine the relationship between the degree of mutant clone selection and the extent of MASLD protection, we deleted *Cideb, Srebf1, Mfn2, Acaca, Mlxipl,* or *Pex5* in the entire liver using high dose AAV-sgRNAs, then treated mice with CDA-HFD. Interestingly, the degree of positive clone selection in CDA-HFD corresponded with the degree of protection from CDA-HFD induced MASLD (**Fig. 3J,K**). Notably, sgRNAs that were not positively selected (*sgAcaca, sgMlxipl, sgPex5*) were associated with modestly beneficial or worse outcomes after CDA-HFD (**Fig. 3J,K**). Thus, different diets can select for different mutations, and the degree of clonal selection correlates with the degree of protection for specific etiologies of liver disease.

### Liver-wide loss of *Cideb* is protective against multiple MAFLD and ALD models

As *Cideb* KO clones were positively selected for in CDA-HFD but not in WD conditions, we asked if *Cideb* suppression might have a greater phenotypic impact in CDA-HFD conditions. To these ends, we generated whole-body and conditional *Cideb* KO mice by using CRISPR-Cas9 germline gene editing. We used sgRNAs to target exon 2 because exons 1 and 2 of *Cideb* do not overlap with *Nop9*, a gene that overlaps in an antisense fashion with the 3’UTR of *Cideb* (**Fig. S5A**). Li’s group previously deleted exon 3-5 of *Cideb* ^3^, which also removed part of the 3’ UTR of *Nop9*. While it is unknown if perturbing *Nop9* has any phenotypic effects, our current KO model avoids this potential off-target effect. DNA and protein analysis confirmed the successful creation of whole-body *Cideb* KO mice missing exon 2 (**Fig. S5B,C**).

Six week old male whole-body *Cideb* WT and KO mice were provided NC, WD, or CDA-HFD for 12 weeks. On NC, there were no differences in body and liver weights between WT and KO mice, but liver/body weight ratios were reduced in KO mice (**Fig. 4A**). Alanine transaminase (ALT) and aspartate transferase (AST) were no different in WT and KO mice, but KO mice showed reduced plasma cholesterol and triglycerides (**Fig. 4B**). On NC, liver histology showed no differences in steatosis between WT and KO mice (**Fig. 4C**). On WD, KO mice had decreased body weight, liver weight, and liver/body weight ratios (**Fig. 4D**). ALT, AST and triglycerides were similar in WT and KO mice, but plasma cholesterol was reduced in the KO mice (**Fig. 4E**). On WD, liver histology showed a reduction in steatosis in KO mice (**Fig. 4C**). On CDA-HFD, KO mice had increased body weight, reduced liver weight, and reduced liver/body weight ratios (**Fig. 4F**). ALT, AST, and plasma cholesterol were decreased in KO mice, but triglycerides were unchanged (**Fig. 4G**). On CDA-HFD, liver histology showed a substantial reduction in steatosis in KO mice (**Fig. 4C**).

**Fig. 4.**
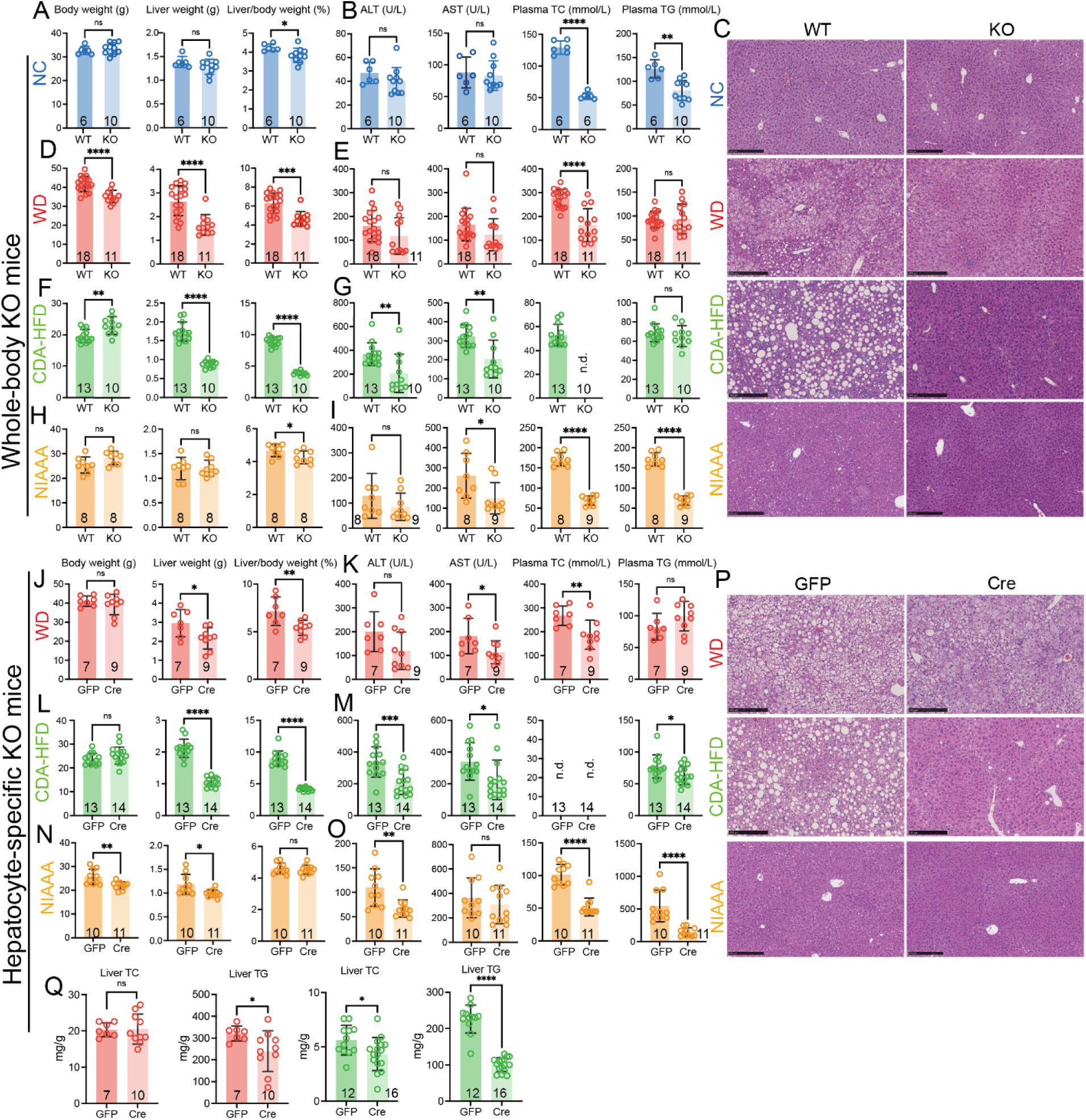
*Cideb* deletion in the liver protects against MAFLD and ALD. **A.** Body weight, liver weight, and liver/body weight ratio of WT and whole-body KO mice fed with NC for 12 weeks. **B.** Plasma ALT, AST, cholesterol, and triglycerides of mice from Fig. 4A. **C.** Representative H&E of livers from mice treated with NC, WC, CDA-HFD, and NIAAA. **D.** Body weight, liver weight, and liver/body weight ratio of WT and whole-body KO mice fed with WD for 12 weeks. **E.** Plasma ALT, AST, cholesterol, and triglycerides of mice from Fig. 4D. **F.** Body weight, liver weight, and liver/body weight ratio of WT and whole-body KO mice fed with CDA-HFD for 12 weeks. **G.** Plasma ALT, AST, cholesterol, and triglycerides of mice from Fig. 4F. **H.** Body weight, liver weight, and liver/body weight ratios of WT and whole-body KO mice fed with the NIAAA diet for 4 weeks with weekly oral gavage of 5 g/kg ethanol. **I.** Plasma ALT, AST, cholesterol, and triglycerides of mice from Fig. 4H. **J.** Body weight, liver weight, and liver/body weight ratio of *Cideb*^fl/fl^ mice injected with AAV-TBG-GFP or AAV-TBG-Cre, fed with WD for 24 weeks. **K.** Plasma ALT, AST, cholesterol, and triglycerides of mice from Fig. 4J. **L.** Body weight, liver weight, and liver/body weight ratios of *Cideb*^fl/fl^ mice injected with AAV-TBG-GFP or AAV-TBG-Cre, fed with CDA-HFD for 12 weeks. **M.** Plasma ALT, AST, cholesterol, and triglycerides of mice from Fig. 4L. **N.** Body weight, liver weight, and liver/body weight ratios of *Cideb*^fl/fl^ mice injected with AAV-TBG-GFP or AAV-TBG-Cre, fed with NIAAA diet for 4 weeks with weekly gavage of 5 g/kg ethanol. **O.** Plasma ALT, AST, cholesterol, and triglycerides of mice from Fig. 4N. **P.** Representative H&E of livers from mice treated with WD, CDA-HFD, and NIAAA. **Q.** Cholesterol and triglyceride content analysis from livers exposed to WD and CDA-HFD. All mice in this figure were male, and all data are presented as mean ± SEM (*p < 0.05; **p < 0.01; ***p < 0.001; ****p < 0.0001).

To determine if *Cideb* has an effect on ALD, we exposed mice to the NIAAA alcohol regimen for 4 weeks (Lieber-DeCarli liquid diet plus weekly oral gavage of ethanol) ^23^. KO mice had reduced liver/body weight ratios (**Fig. 4H**), as well as reduced plasma AST, cholesterol, and triglycerides (**Fig. 4I**). On NIAAA, liver histology showed a modest reduction in steatosis in KO mice (**Fig. 4C**). Overall, whole-body *Cideb* deletion protected mice from the effects of WD, CDA-HFD, and alcohol, but most effectively mitigated the effects of CDA-HFD (**Fig. 4C**).

To determine if organ-specific loss of *Cideb* mimics germline deletion, we also generated *Cideb* floxed mice with loxP sequences flanking exon 2 (**Fig. S5B**). We used AAV-TBG-Cre to delete *Cideb* in hepatocytes at 6 weeks of age, resulting in the absence of CIDEB protein in the liver (**Fig. S5D**). *Cideb* WT and hepatocyte-specific KO mice were provided with a WD for 24 weeks or CDA-HFD for 12 weeks. After 24 weeks of WD feeding, mice with hepatocyte-specific deletion had decreased liver weight and liver/body weight ratios (**Fig. 4J**), but in contrast to the whole-body KO model, the hepatocyte-specific KO did not cause body weight reduction. Plasma AST and cholesterol levels were reduced in the hepatocyte-specific KO, while ALT trended down and triglyceride was unchanged (**Fig. 4K**). On CDA-HFD, hepatocyte-specific KO mice had similar body weights, but lower liver, and liver/body weight ratios (**Fig. 4L**). Plasma ALT, AST, and triglycerides were lower in the hepatocyte-specific KO mice and plasma cholesterol was below the limit of detection for both WT and KO mice (**Fig. 4M)**. Hepatocyte-specific KO mice exposed to the NIAAA diet showed reduced body and liver weights, but the liver/body weight ratios did not change (**Fig. 4N**). Similar to the whole-body KO mice, plasma ALT, triglyceride, and cholesterol were reduced, but AST remained unchanged (**Fig. 4O**). In all MASLD and ALD models, hepatocyte-specific KO mice had reduced steatosis on histology; the magnitude of decrease was most pronounced with CDA-HFD (**Fig. 4P**). KO livers on both MASLD diets showed reduced liver triglycerides and KO livers on CDA-HFD showed reduced liver cholesterol (**Fig. 4Q**). For both diets, replicate experiments in female hepatocyte-specific KO mice showed similar but less pronounced phenotypes (**Fig. S6**). Altogether, both whole-body and hepatocyte-specific deletion of *Cideb* impaired the development of MASLD and ALD. The hepatocyte-specific KO mice did not show weight loss, thus metabolic improvements were not simply a consequence of body weight change.

### After MAFLD establishment, hepatocyte-specific *Cideb* deletion can reverse disease

We performed genetic experiments to mimic a potential hepatocyte targeting siRNA approach for MAFLD. We asked if hepatocyte-specific *Cideb* deletion after the establishment of MAFLD could reverse some features of MAFLD. We used two cohorts of mice for this experiment. The first cohort, referred to as the ‘Prevention group’, was discussed in the previous section. These male *Cideb^fl/fl^*mice were given AAV-TBG-GFP (control) or AAV-TBG-Cre to induce *Cideb* deletion, then were given CDA-HFD for 12 weeks (**Fig. 5A**). The second cohort, referred to as the ‘Treatment group’, consisted of male *Cideb^fl/fl^*mice that were fed CDA-HFD for 6 weeks, then given AAV-TBG-GFP or AAV-TBG-Cre, then resumed on CDA-HFD for an additional 6 weeks (**Fig. 5A**). In the ‘Prevention group’, MASLD characteristics were improved as discussed above (**Fig. 4L,M,P**) and fibrosis was decreased (**Fig. 5B**). In the ‘Treatment group’, KO mice had similar body weights, but reduced liver weights and liver/body weight ratios (**Fig. 5C**). Plasma ALT, AST, and triglyceride levels were similar in WT and KO mice (**Fig. 5D**). In the ‘Treatment’ experiment, control mice had similar levels of steatosis between 6 and 12 weeks, but KO mice had a robust reversal of steatosis after 6 weeks of CDA-HFD (**Fig. 5E**). That said, the modest amount of fibrosis observed was unchanged between WT and KO groups (**Fig. 5F**). Many of the same phenotypic patterns were reproduced in female mice (‘Prevention’ in **Fig. S6** and ‘Treatment’ in **Fig. 5G-J**). This suggested that hepatocyte-specific deletion of *Cideb* was sufficient to reverse some, but not all features of MASH.

**Fig. 5.**
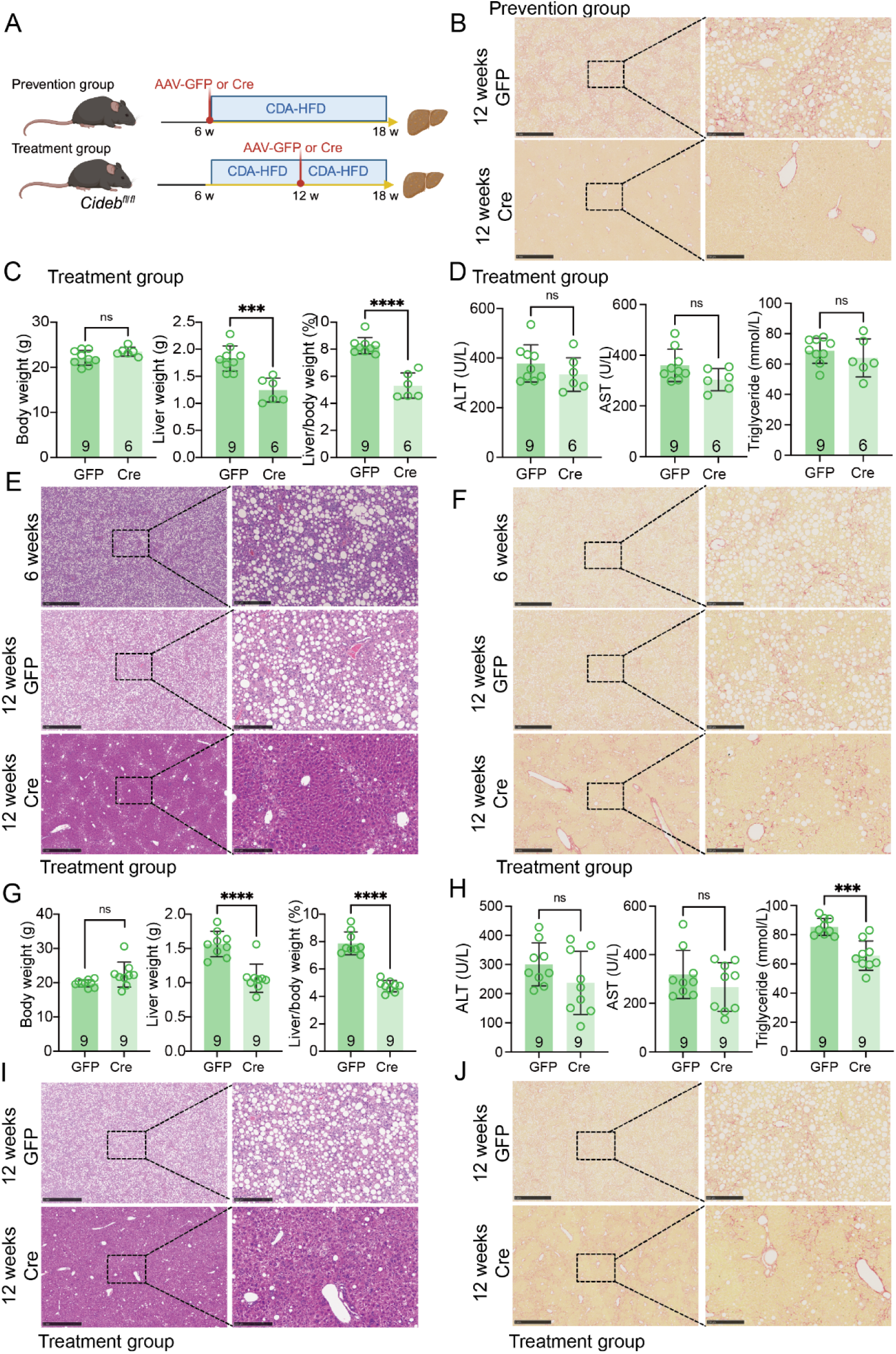
*Cideb* deletion reverses MAFLD progression in male and female mice. **A.** Experimental schema for the “Prevention” and “Treatment” groups under CDA-HFD. In the “Prevention” group, *Cideb^fl/fl^* mice were injected with AAV-TBG-GFP or AAV-TBG-Cre at 6 weeks of age and fed CDA-HFD for 12 weeks (data shown in Fig. 4L). In the “Treatment” group, *Cideb^fl/fl^* mice were fed CDA-HFD starting at 6 weeks of age for 6 weeks, then given AAV-TBG-GFP or AAV-TBG-Cre, then maintained on CDA-HFD for another 6 weeks before harvest. **B.** Representative Sirius red staining of male “Prevention” group livers from Fig. 4L. **C.** Body weight, liver weight, and liver/body weight ratio of male “Treatment” mice. **D.** Plasma ALT, AST, and triglycerides from the male “Treatment” mice. **E.** Representative H&E staining of *Cideb^fl/fl^* livers from the 6 week timepoint (when the AAV was given), and from the control and KO male “Treatment” mice at the 12 week timepoint. **F.** Representative Sirius red staining of livers from the male “Treatment” mice. **G.** Body weight, liver weight, and liver/body weight ratio of female “Treatment” mice. **H.** Plasma ALT, AST, and triglycerides from the female “Treatment” mice. **I.** Representative H&E staining of livers from the female “Treatment” mice. **J.** Representative Sirius red staining of livers from the female “Treatment” mice. All data are presented as mean ± SEM (***p < 0.001; ****p < 0.0001).

### Unsaturated fats preferentially accumulate in LDs and *Cideb* deletion mitigates this

The impact of *Cideb* deletion in the CDA-HFD setting was increased in comparison to the WD setting. Lipidomics revealed differences in the types of free fatty acids (FFAs) present in the diets. The proportion of unsaturated fatty acids was higher in CDA-HFD than in WD (**Fig. S7A**). To understand if differences in the diets were also seen within the livers of mice, we performed FFA analysis on livers exposed to WD or CDA-HFD. CDA-HFD fed livers contained a higher proportion of unsaturated vs. saturated fatty acids compared to WD fed livers (**Fig. S7B**).

To determine if unsaturated fats accumulate in LDs more than saturated fats ^24,25^, we provided different saturated and unsaturated fatty acids (100 µM, 16h) to human (HepG2, Huh7) and mouse (H2.35) liver cancer cell lines. Saturated fatty acids included palmitic (16:0), stearic (18:0), and arachidic acid (20:0). The unsaturated fatty acids included oleic (18:1), linolenic (18:3), stearidonic (18:4), mead (20:3), arachidonic (20:4), and eicosapentaenoic acid (20:5). Compared to saturated fatty acids, the unsaturated fatty acids resulted in the formation of larger LDs (**Fig. S7C,D**). Given the potential differences between benign and malignant liver cells, we repeated these experiments using primary hepatocytes from C57BL/6 mouse livers. The results were consistent with the cancer cell lines (**Fig. S7E,F**), supporting the idea that unsaturated fatty acids form LDs more efficiently than saturated fatty acids in the liver.

To determine if *Cideb* KO mice have altered accumulation of fatty acid species, we performed FFA analysis on WT and *Cideb* KO liver samples from mice fed the CDA-HFD. There was little change in the relative levels of saturated fatty acids between WT and KO, but unsaturated fatty acids were disproportionately decreased in KO livers (**Fig. S8A**). To corroborate these findings, we also incubated primary hepatocytes from control *Cideb^fl/fl^* and liver-specific *Alb-Cre; Cideb^fl/fl^* mice in media containing predominantly saturated or unsaturated fatty acids. While unsaturated fatty acids resulted in larger LDs in WT hepatocytes, saturated fatty acids did not have a similar effect on LD formation (**Fig. S7C,F**). However, in KO hepatocytes, unsaturated fatty acids were unable to form LDs (**Fig. S8B,C**). Overall, LDs were smaller in KO hepatocytes, regardless of exogenous fatty acid source. This suggests that deleting *Cideb* impairs the enlargement of LDs mainly through the reduction of unsaturated fatty acids.

### Loss of *Cideb* did not reduce DNL or increase VLDL secretion

While CDA-HFD results in more unsaturated fatty acid accumulation within LDs, and *Cideb* deletion primarily reduces accumulation of unsaturated fatty acids, it is still unclear why there is a net reduction of neutral lipids in KO livers. We sought to determine if *Cideb* deficiency alters lipid delivery/uptake, synthesis, secretion, or oxidation. MAFLD is most commonly associated with obesity. As indicated previously, whole-body *Cideb* deletion resulted in an increase in body weight in mice fed the CDA-HFD (**Fig. 4F**), so body weight cannot explain the reduced steatosis seen in the *Cideb* deficient mice. The increased *Cideb* KO body weight could be caused by reduced *Gdf15* and/or *Fgf21* secondary to improved liver health (**Fig. 8D**).

We then assessed DNL pathways to determine if decreased lipogenesis could account for improved MASLD ^26^. KO livers from mice fed WD showed increases in *Chrebp*, *Dgat1* and *Dgat2*, but no changes in the other DNL mRNAs (**Fig. 6A**). KO livers from mice fed CDA-HFD showed an increase in DNL pathway mRNAs including *Srebp1c*, *Chrebp*, *Acly*, *Acc1*, *Acacb*, *Fasn*, *Gpat1*, *Dgat1*, *Dgat2* and *Pparγ* (**Fig. 6B**). In addition, SREBF1 and HMGCR were lower in KO livers on WD (**Fig. 6C**), but SREBF1, SREBF2, FASN, HMGCR, ACC1 and PPARγ protein expression were higher in KO livers on CDA-HFD (**Fig. 6C**). This suggests that on CDA-HFD, DNL is likely to be increased rather than decreased in KO livers, whereas on WD, DNL was only modestly lower in KO livers.

**Fig. 6.**
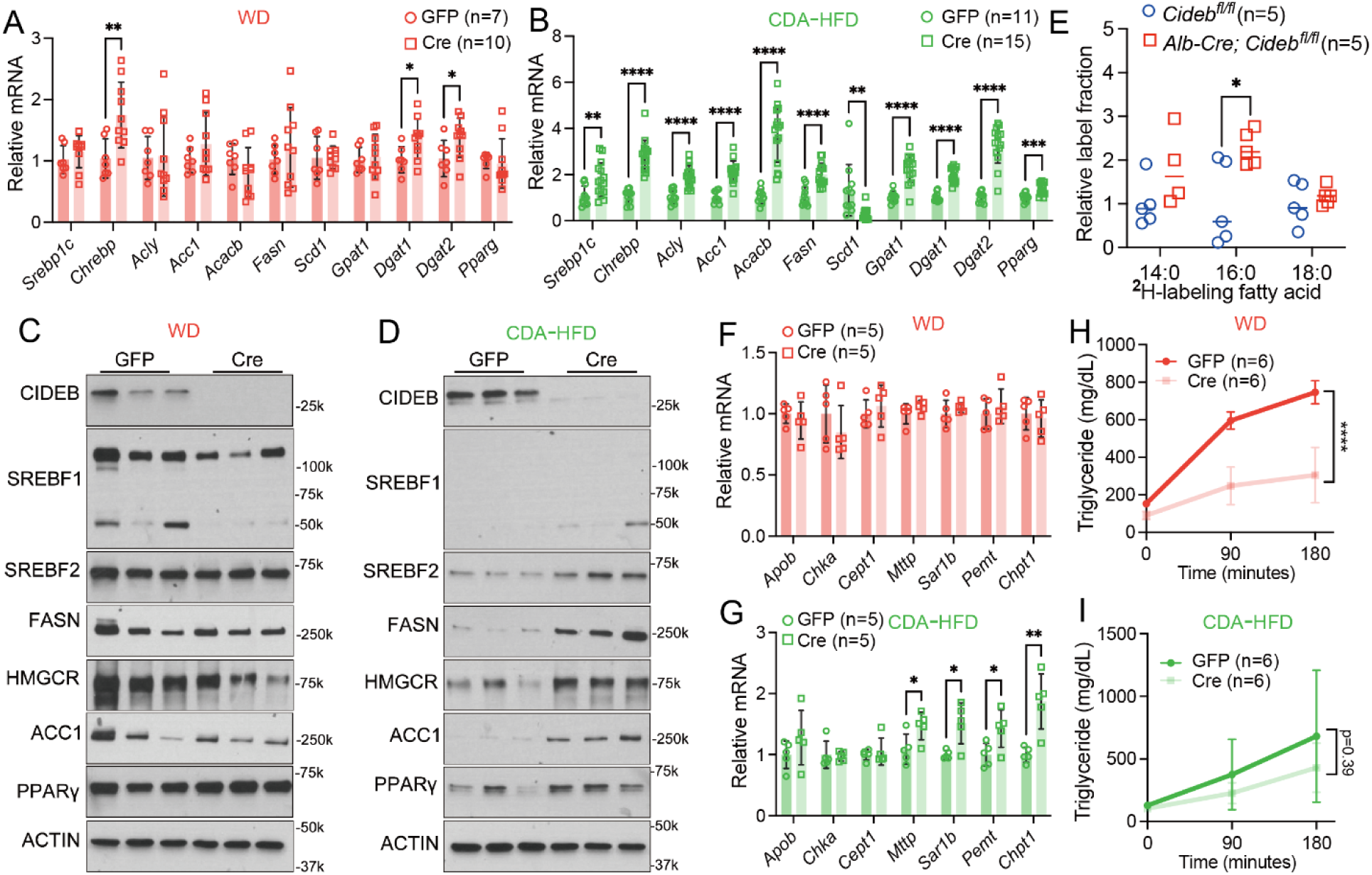
Influence of *Cideb* KO on DNL and VLDL lipidation/secretion. **A.** qPCR of DNL genes from livers fed with 24 weeks of WD. **B.** qPCR of DNL genes from livers fed with 12 weeks of CDA-HFD. **C.** Western blot analysis of DNL-related proteins in livers exposed to WD for 24 weeks. **D.** Western blot analysis of DNL-related proteins in livers exposed to CDA-HFD for 12 weeks. **E.** Total labeled fraction of liver fatty acid species after 5 hours of short-term ^2^H_2_O tracing in *Cideb^fl/fl^* and *Alb-Cre; Cideb^fl/fl^* mice. **F.** VLDL-related mRNA expression from livers treated with 24 weeks of WD. Data taken from RNA-seq analysis. **G.** VLDL-related mRNA expression from livers treated with 12 weeks of CDA-HFD. Data taken from RNA-seq analysis. **H.** Poloxamer-407 assay on mice fed with WD for 1 week and fasted overnight. Plasma triglyceride levels were measured. **I.** Poloxamer-407 assay on mice fed with CDA-HFD for 1 week and fasted overnight. All data are presented as mean ± SEM (*p < 0.05; **p < 0.01; ***p < 0.001; ****p < 0.0001).

To functionally assess DNL in the CDA-HFD setting, we performed isotope enrichment analysis with a single dose of deuterated water (²H₂O) followed by assessment of liver samples 5 hours later. In this assay, ²H₂O effectively labels the hydrogen atoms in fatty acids via transfer from NADPH during fatty acid synthesis ^27,28^. Therefore, ²H₂O is used to measure DNL by quantifying the relative ²H enrichment of lipids ^27,28^. While labeled fatty acids between WT and KO livers fed CDA-HFD were similar, palmitic acid was increased in the KO suggesting modestly increased DNL (**Fig. 6E**). Thus, *Cideb* deficient mice were not protected from MASLD through decreases in DNL.

Increased disposal of lipids through very low density lipoprotein (VLDL) lipidation/secretion could represent an alternative mechanism for reduced MAFLD. However, *Cideb* deletion is predicted to impair VLDL lipidation based on prior reports ^29^ and CDA-HFD induces steatosis at least in part by impairing VLDL secretion as is reflected in the low plasma triglycerides seen in this context. Nevertheless, we considered this pathway in our models. In the RNA-seq data from the WD model, the expression of key VLDL-related mRNAs was similar in WT and KO livers (**Fig. 6F**). In the RNA-seq data from the CDA-HFD model, *Apob*, *Chka*, and *Cept1* were similar in WT and KO livers, while *Mttp*, *Sar1b*, *Pemt*, and *Chpt1* were modestly increased in KO livers (**Fig. 6G**). Next, we functionally tested VLDL secretion using the poloxamer 407 assay. This polymer inhibits Lipoprotein lipase, thereby reducing hydrolysis of triglycerides and facilitating VLDL accumulation. In this assay, we observed reduced VLDL secretion in KO mice on both WD and CDA-HFD (**Fig. 6H,I**), which is consistent with *Cideb* KO mice fed WD having reduced plasma cholesterol (**Fig. 4E,K**), an observation that is also favorable for therapeutic approaches. Overall, these data suggest that *Cideb* deletion does not protect from MASLD through increased lipid export from the liver.

### *Cideb* deletion in the liver in mice increases hepatic FAO

The CDA-HFD RNA-seq Gene Set Enrichment Analysis (GSEA) revealed upregulation of pathways related to bile acid metabolism, fatty acid metabolism, xenobiotic metabolism, adipogenesis, oxidative phosphorylation, and peroxisomes in KO vs. WT livers (**Fig. S9A,B**). In the context of both diets, several individual mRNAs related to FAO (*Acads, Acsf2, Acsm1, Cpt1a, Cpt1b, Cpt2, Acadl1, Acadv1, Acadm, Ppara*) and peroxisomes (*Agps, Pex14, Pex16, Pex5, Acaala, Acaalb, Pex11a, Agt, Cat, Ehhadh, Acox1, Acox2, Acsl1*) were upregulated (**Fig. 7A-D**). Proteins involved in FAO and peroxisome pathways were also increased in KO livers, but more in the CDA-HFD than in the WD setting (**Fig. 7E,F**).

**Fig. 7.**
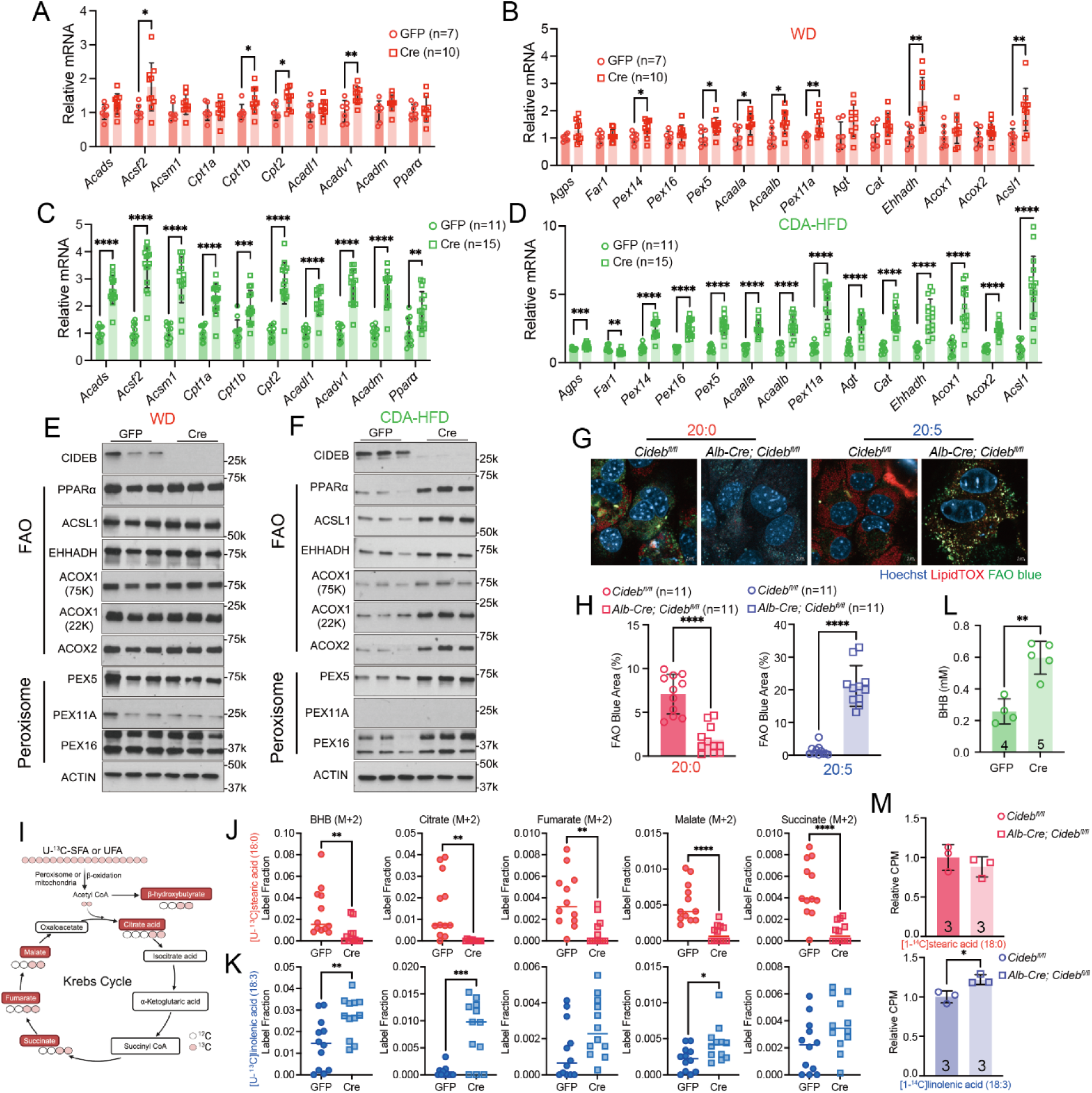
*Cideb* deletion increases FAO. **A.** qPCR of FAO genes from livers fed with 24 weeks of WD. **B.** qPCR of peroxisome-related genes from livers fed with 24 weeks of WD. **C.** qPCR of FAO genes from livers fed with 12 weeks of CDA-HFD. **D.** qPCR of peroxisome-related genes from livers fed with 12 weeks of CDA-HFD. **E.** Western blot analysis of FAO and peroxisome-related proteins in livers exposed to WD for 24 weeks. The CIDEB and ACTIN blots are the same as shown in Fig. 6C. **F.** Western blot analysis of FAO- and peroxisome-related proteins in livers exposed to CDA-HFD for 12 weeks. The CIDEB and ACTIN blots are the same as shown in Fig. 6D. **G.** Representative FAO blue (green) of primary hepatocytes treated with arachidic acid (20:0) or arachidonic acid (20:5) overnight from *Cideb*^fl/fl^ and *Alb-Cre; Cideb^fl/fl^* mice. LDs were stained with LipidTOX (red) and nuclei were stained with Hoechst (blue) (scale bars = 5 µm). **H.** Quantification of LD area from Fig. 7G. **I.** Schematic of [U-^13^C]linolenic and [U-^13^C]stearic acid tracing. **J.** Fractional enrichment of indicated liver isotopologues in *Cideb^fl/fl^* and *Alb-Cre; Cideb^fl/fl^* mice after 2-hour infusion with [U-^13^C]stearic acid (n=12, 12 mice). **K.** Fractional enrichment of indicated liver isotopologues in *Cideb*^fl/fl^ and *Alb-Cre*; *Cideb*^fl/fl^ mice after 2-hour infusion with [U-^13^C]linolenic acid (n=12, 12 mice). **L.** Plasma BHB from *Cideb^fl/fl^* mice injected with AAV-TBG-GFP or AAV-TBG-Cre fed with CDA-HFD for 1 month. **M.** Relative FAO activity was assessed by measuring counts per minute (CPM) of ^14^CO_2_ production in primary hepatocytes isolated from *Cideb*^fl/fl^ and *Alb-Cre*; *Cideb*^fl/fl^ mice. The hepatocytes were treated with [1-^14^C]stearic acid or [1-^14^C]linolenic acid to evaluate the effect of *Cideb* deletion on fatty acid metabolism. All data are presented as mean ± SEM (*p < 0.05; **p < 0.01; ***p < 0.001; ****p < 0.0001).

**Fig. 8.**
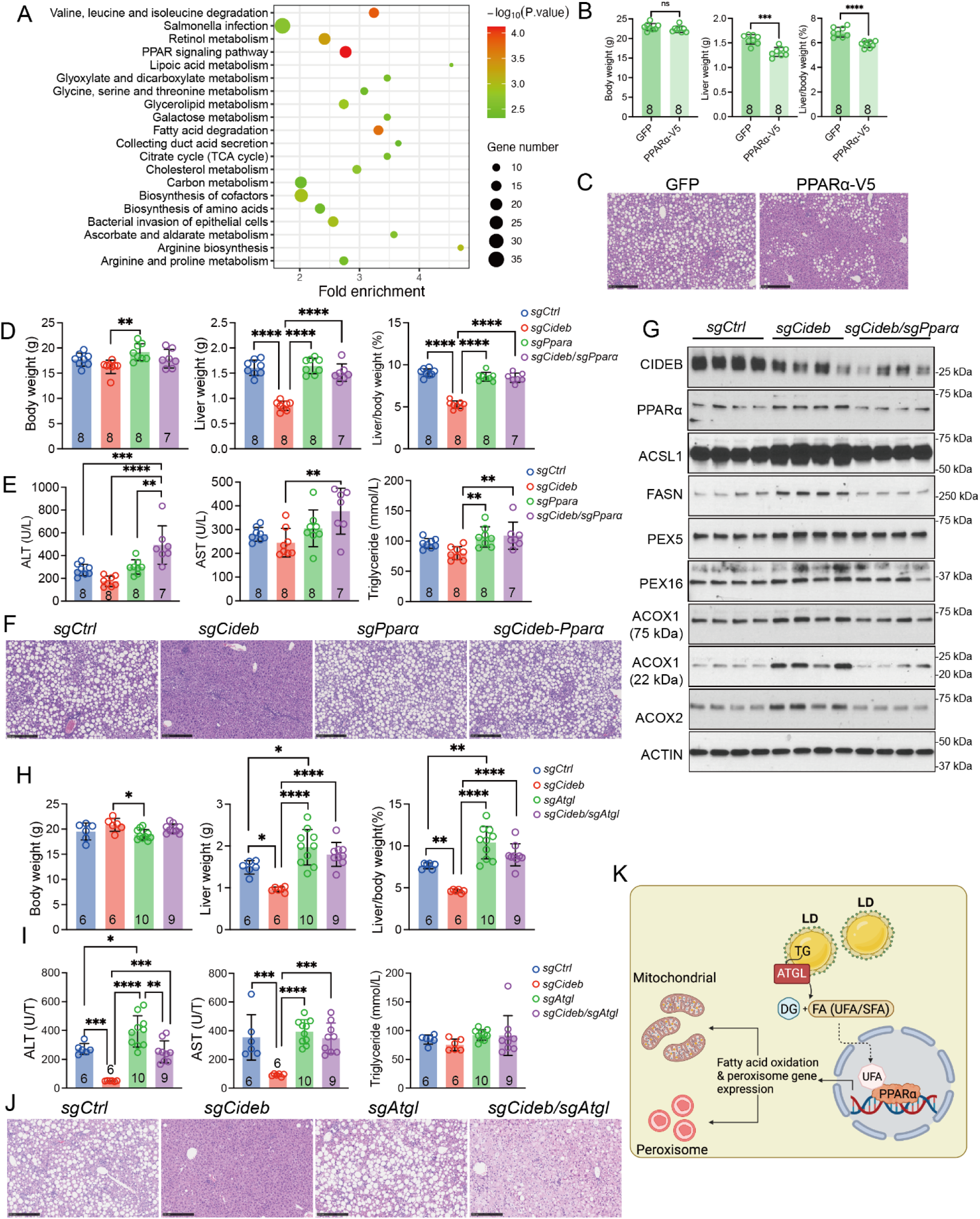
*Pparα* and *Atgl* deletion rescue the protective effects of *Cideb* deletion. **A.** Bubble plot showing differentially expressed pathways between WT and KO livers treated with 12 weeks of CDA-HFD. **B.** Body weight, liver weight, and liver/body weight ratio of WT mice injected with AAV-TBG-GFP or AAV-TBG-PPARα-V5 and treated with CDA-HFD for 3 weeks. **C.** Representative H&E staining of livers from Fig. 8B. **D.** Body weight, liver weight, and liver/body weight ratio in male iCas9 mice given high-dose AAV-sgRNAs (*sgCtrl*, *sgCideb*, *sgPparα*, or *sgCideb*/*sgPparα*). At 8.5 weeks, mice were fed CDA-HFD for 3 weeks. **E.** Liver function tests from mice in Fig. 8D. **F.** Representative H&E staining of livers from Fig. 8D. **G.** Western blot analysis of livers from Fig. 8D. **H.** Body weight, liver weight, and liver/body weight ratio in male iCas9 mice given high-dose AAV-sgRNAs (*sgCtrl*, *sgCideb*, *sgAtgl*, or *sgCideb*/*sgAtgl*). At 8.5 weeks, mice were fed CDA-HFD for 3 weeks. **I.** Liver function tests from mice in Fig. 8H. **J.** Representative H&E staining of livers from Fig. 8H. **K.** Model for how ATGL affects LDs, which subsequently influences PPARα function. All data are presented as mean ± SEM (*p < 0.05; **p < 0.01; ***p < 0.001; ****p < 0.0001).

To quantify FAO in primary hepatocytes with and without CIDEB, we used a reagent that visualizes FAO with green fluorescence. In cells exposed to the saturated arachidic acid (20:0), FAO blue signal was reduced in *Cideb* KO vs. WT hepatocytes, while in cells exposed to the unsaturated arachidonic acid (20:4), FAO blue signal was increased in KO vs. WT hepatocytes (**Fig. 7G,H**). To evaluate in vivo changes in FAO, *Cideb*^fl/fl^ and *Alb-Cre*; *Cideb*^fl/fl^ mice were infused with unsaturated ([U-^13^C]linolenic acid (18:3)) or saturated fatty acids ([U-^13^C]stearic acid (18:0)) for 2 hours (**Fig. 7I**). In KO mice, injection with saturated fatty acids led to a significant decrease in ^13^C-labeled β-hydroxybutyrate (BHB), citrate, fumarate, malate, and succinate, whereas injection with unsaturated fatty acids resulted in an increase in these metabolites, indicating increased FAO (**Fig. 7J,K**). To assess the overall production of ketones through FAO, we measured BHB in the plasma of CDA-HFD treated mice, showing that liver-specific KO mice had higher plasma BHB levels (**Fig. 7L**). Consistently, similar findings were made in primary hepatocytes using a gold standard radioactive assay that directly measures CO₂ release from fatty acids. In KO hepatocytes, this showed increased β-oxidation when linoleic acid is provided, but no significant difference with stearic acid (**Fig. 7M**). Collectively, these results suggest a relative increase in β-oxidation of unsaturated fatty acids, and an overall increase in FAO in *Cideb* KO mice.

### PPARα and ATGL are effectors of increased FAO in *Cideb* KO livers

KEGG pathway analysis indicated that PPAR signaling was upregulated on the CDA-HFD (**Fig. 8A**). Specifically, peroxisome proliferator-activated receptor alpha (PPARα) was upregulated at both the mRNA and protein levels (**Fig. 7C,F**). Given that PPARα is a nuclear receptor and transcription factor that regulates lipid catabolism and has been implicated in mediating increased FAO in the context of altered LD fatty acid regulation ^30–32^, we hypothesized that upregulation of PPARα might be a key determinant of the increased β-oxidation seen in *Cideb* KO livers. We first asked if PPARα overexpression alone is enough to reduce fatty liver disease. AAV was used to overexpress GFP or PPARα-V5 in WT mice, then mice were fed with CDA-HFD. Three weeks later, livers with PPARα overexpression showed reduced steatosis (**Fig. 8B,C**).

While overexpression of PPARα alone is sufficient to inhibit MAFLD, it is unclear if PPARα is required for the protective phenotype caused by *Cideb* loss. We used iCas9 mice and AAV-sgRNA vectors to generate control, *Cideb* single, *Pparα* single, or *Cideb*; *Pparα* double KO livers in the context of 3 weeks of CDA-HFD (**Fig. S9C**). The liver/body weight ratios were similar in control, *Pparα* KO, and *Cideb*; *Pparα* double KO mice, but all were greater than in *Cideb* single KO mice (**Fig. 8D**). Double KO mice had higher ALT, AST and triglyceride levels than control and single KO mice (**Fig. 8E**). Similarly, H&E staining showed extensive steatosis in control, *Pparα* KO, and *Cideb*; *Pparα* double KO mice, and all of these groups had substantially more steatosis than *Cideb* KO mice (**Fig. 8F**). CIDEB and PPARa were successfully reduced at the protein level, and *PPARα* deletion groups rescued or inhibited the downstream peroxisome associated protein increases caused by *Cideb* loss (**Fig. 8G**). This suggests that deleting *Cideb* upregulates the transcriptional activity of PPARα, promoting the expression of peroxisome- and FAO-related genes.

However, it was still unclear how loss of *Cideb* stimulated increased PPARα levels and transcriptional activity that promoted β-oxidation. We hypothesized that in *Cideb* KO livers, LD triglycerides undergo lipolysis, resulting in FFAs that act as ligands to activate PPARα. To test this, we asked if the inhibition of ATGL-mediated triglyceride lipolysis in hepatocytes could block the effects of *Cideb* deletion ^33,34^. To do this, we also generated control, *Cideb* single, *Atgl* single, or *Cideb*; *Atgl* double KO livers in the context of 3 weeks of CDA-HFD (**Fig. S9C**). The liver/body weight ratios were similar in control, *Atgl* KO, and *Cideb*; *Atgl* double KO mice. All of these groups had greater LW/BW ratios than *Cideb* single KO mice (**Fig. 8H**). Similarly, H&E staining showed extensive steatosis in control, *Atgl* KO, and *Cideb*; *Atgl* double KO mice, whereas *Cideb* single KO mice had minimal steatosis (**Fig. 8I-J**). Single *Atgl* KO livers showed the most steatosis. In *Cideb*; *Atgl* double KO mice, LDs were smaller but did not disappear as seen with single *Cideb* KO (**Fig. 8I**). This suggests that ATGL-mediated lipolysis contributes to the activation of PPARa in *Cideb* KO livers (**Fig. 8K**).

### *Cideb* loss prevents liver cancer development in the context of MAFLD

Hepatocellular carcinoma is one of the most devastating outcomes associated with MASH ^35^. Given that *Cideb* loss can promote clone expansion in MASH, we wanted to ask if this might eventually lead to increased cancer development. We used the diethylnitrosamine (DEN) plus WD approach to instigate tumor growth. 14-day-old *Cideb^fl/fl^* mice were injected intraperitoneally with DEN. At 28 days of age, these mice were injected with either AAV-TBG-GFP or AAV-TBG-Cre and initiated on WD. After 8 months, more HCC was found on the liver surface of WT vs. hepatocyte-specific KO male mice (**Fig. S10A**), while there was only a non-significant difference in female mice (**Fig. S10A**). Microscopic tumor numbers were also lower in male KO livers (**Fig. S10B,C**).

## Discussion

Somatic and germline *CIDEB* mutations appear to ameliorate liver disease in humans ^1,2^. Here, functional characterization of all reported *CIDEB* somatic mutations showed that they impair CIDEB’s impact on LD growth. Furthermore, mosaic deletion of *Cideb* in the murine liver leads to positive selection of deleted clones, but specifically in the setting of particular fatty liver inducing diets. Interestingly, *Cideb* somatic mutations were more strongly selected for in the context of a CDA-HFD compared with a WD diet. In keeping with these data, liver-specific *Cideb* deletion was associated with greater protection against liver disease associated with CDA-HFD feeding than WD feeding. This suggests that *Cideb* directed interventions could be more effective in some subtypes of fatty liver disease. Future clinical trials in MASH may benefit from considering different mechanisms of MASH development, or by exploiting *Cideb* somatic mutations as biomarkers to predict the patients that would benefit the most.

Elegant studies from Peng Li’s group have shown that whole-body *Cideb* deletion is protective against fatty liver disease and hyperlipidemia in mouse models ^3,7^. While they showed that CIDEB promotes VLDL lipidation and restrains β-oxidation ^3^, the predominant mechanism for MASLD protection was related to DNL. They showed that CIDEB promotes the loading of SREBP/SCAP into COPII vesicles, thereby facilitating SREBP1 transit from ER to nucleus ^5^. In this way, CIDEB promotes DNL transcription and its deletion impairs lipogenesis in the context of high fructose, low fat diets. While this study also shows anti-MAFLD phenotypes, our results suggest that altered β-oxidation, rather than DNL, is predominantly responsible for protective phenotypes. The differences in the importance of DNL could be due to differences in the MAFLD inducing diets used between studies, together with the fact that all other studies were only done in whole-body *Cideb* KO mice, which also manifest a reduction in body weight. Since there are several therapeutic approaches associated with DNL inhibition in clinical testing, it is important to delineate alternative approaches that are focused on β-oxidation.

Besides demonstrating that loss of function mechanisms underscore positive clonal selection in fatty liver disease, our studies also increase the mechanistic understanding of CIDEB function. *Cideb* KO hepatocytes have smaller LDs, an observation that is associated with reduced neutral lipids and cholesterol storage in the liver. In addition, there were significant decreases in unsaturated fatty acid content in the livers of *Cideb* KO mice fed with CDA-HFD, while saturated fatty acids remained largely unaffected. This suggests that CIDEB preferentially regulates the metabolism, storage, and FAO of unsaturated fatty acids in hepatocytes. *Cideb* loss reduces neutral lipid storage in LDs not through reduced DNL, but rather through increased lipolysis and β-oxidation. Since ATGL is the main triglyceride lipase in the liver, and because *Atgl* deletion rescues the effects of *Cideb* deletion, the reduction in LD size is potentially mediated through ATGL. Alternatively, the reduction in LD size caused by *Cideb* deletion could increase the surface area to volume ratio of LDs, thus increasing the efficiency of lipolysis by ATGL. Since unsaturated fatty acids can serve as ligands that activate PPARα’s transcriptional activity ^36–38^, we reason that increased unsaturated FFAs levels trigger increased β-oxidation through PPARα. Consequently, the enhanced expression of PPARα-regulated pathways facilitates the metabolism and consumption of FFAs in hepatocytes, thereby protecting the liver from the detrimental effects of unsaturated fatty acid accumulation.

In conclusion, our findings show that targeting CIDEB may offer potential therapeutic strategies for mitigating MAFLD and related metabolic disorders characterized by aberrant lipid metabolism. While CIDEB operates at multiple levels of lipid metabolism, our work suggests that a disease defining manifestation of *Cideb* mutation is increased β-oxidation.

## Acknowledgements

We would like to thank Liang Dong and Anna Szczoczarz for assisting in the cloning of human *CIDEB*. We also thank J. Shelton (UTSW histopathology Core), E. Nwoka, and C. Moxon (UT Southwestern Tissue Management Shared Resource) for histopathology; D. Ramirez (UTSW Whole Brain Microscopy Facility, RRID: SCR_017949) for whole-liver imaging; S. Wu, Y. J. Kim, and J. Lyu (CRI Sequencing Core) for sequencing; and S. Hacker, A. Walker, and J.I. Gamayot for metabolic phenotyping assays. D.B.S. is supported by the Wellcome Trust (WT 219417), the MRC (MR/X00970X/1), and The National Institute for Health Research (NIHR) Cambridge Biomedical Research Centre and NIHR Rare Disease Translational Research Collaboration. This work was supported by the Tissue and Cell Imaging Facility at the Institute of Metabolic Science, who were funded by the Medical Research Council (MC_UU_00039). H.Z. is supported by the Moody Foundation, the NIH (R01AA028791, R01CA251928, DP1DK139976), and a Simmons Comprehensive Cancer Center Cancer and Obesity Translational Pilot Award.

## Author Contributions

Q.Z., S.P., D.S. and H.Z. conceived the project, performed the experiments, and wrote the manuscript. X.W. and P.M. performed lipidomics and isotopologue analysis. M-H.H. performed bioinformatic analysis on the MOSAIC screens. Z.L. and X.F. performed immunofluorescence and Sirius red staining. X.R. performed primary hepatocyte isolation. S.L measured liver lipids. J.W., L.L., A.B., M.W. and X.G. performed animal work. G.Mannino performed the poloxamer assay. G.Maggiore, M.Z. and B.L. performed RNA-seq analysis. S.P., K.L., Z.Q., P.P. and M.O.M, performed the human CIDEB and CIDEC experiments. D.K. and G.H. performed isotope tracing and analysis for FAO studies. T.S. and Y.Z. generated the *Cideb* KO mice. V.S carried out the structural modeling analysis for human CIDEB and CIDEC. P.G. performed the histologic analysis and NAS scoring. M.H. and P.C. provided somatic mutation data.

## Declaration of Conflicts

H.Z. and P.C. are co-founders of Quotient Therapeutics and Jumble Therapeutics, is an advisor for Newlimit, Alnylam Pharmaceuticals, and Chroma Medicines. H.Z. receives research support from Chroma Medicines and owns stock in Ionis and Madrigal Pharmaceuticals. H.Z. and L.L. have a patent on CIDEB siRNA for liver disease (patent #63/328,557).

## Materials and Methods

### Cloning

Human *CIDEA*, *CIDEB,* and *CIDEC* cDNAs were amplified by PCR using Phusion High-Fidelity DNA Polymerase and cloned into pCDNA3.1 either untagged or tagged with HA at the C-termini (Thermo Fisher, F530L). Truncations (stop), point mutants, and deletions (Δ) were generated using either QuikchangeII XL (Agilent Technologies) site-directed mutagenesis or Gibson assembly. WT and mutant versions of *CIDEA*, *CIDEB,* and *CIDEC* cDNAs were also inserted into a pEGFPN3 vector (Clontech, 6080-1) to create C-terminal GFP tagged constructs.

### Cell lines

Huh7, HepG2, and HeLa cells (ATCC) were cultured in High glucose Dulbecco’s Modified Eagle’s Medium (DMEM) supplemented with 10% (vol/vol) fetal bovine serum (FBS), and 1% penicillin-streptomycin at 37°C in 20% O_2_ and 5% CO_2_. H2.35 cells were cultured with complete DMEM supplemented with 4% FBS, 200 nM dexamethasone, and 1x penicillin-streptomycin.

### High content confocal imaging and quantification of LD size

HeLa cells were seeded at a density of 1×10^5^ cells/well onto ethanol-treated glass coverslips in 12-well plates or 6000 cells/well onto Perkin Elmer 96-Well Phenoplate Ultra Plates (6055302; Perkin Elmer). The following day, *CIDEA*, *CIDEB,* or *CIDEC* cDNA was co-transfected along with GFP using Lipofectamine LTX according to the manufacturer’s instructions. 4 h later, the media was replaced with fresh DMEM supplemented with 400 μM Oleic acid. 24 h post-transfection, the media was removed and cells were washed three times with 1x PBS and fixed with 4% (v/v) formaldehyde in PBS for 15 min at room temperature followed by three washes for 5 min with PBS. LDs were visualized either by addition of 1:2000 dilution of BODIPY™ 558/568 C_12_ during oleate-loading or stained with 1x HCS LipidTOX Deep Red Neutral Lipid Stain post-fixation for one hour. Imaging was carried out on the Perkin Elmer Opera Phenix spinning disc confocal microscope equipped with a 40x/1.1 Water objective with 2-pixel binning. EGFP and LipidTox Deep Red Neutral Stain were excited at 488 and 640, and emission signals were collected at 500-550 nm and 650-760 nm, respectively. Images were acquired and LD volumes in GFP positive cells were determined by the Harmony phenoLOGIC Software (v4.9; Perkin Elmer). LD enlargement activity is represented as Z-scores where data is normalised as *x-μ/σ* where *μ* was calculated from wild type (WT) CIDE or empty vector (EV) and *σ* from all values.

For HA immunostaining, fixed cells were permeabilized with 0.5% saponin for 10 min, washed with PBS, blocked with 1% BSA in PBST for 1h, followed by overnight incubation at 4C with 1:500 HA-Tag (C29F4) antibody. Cells were washed four times for 5min with 0.1% BSA in PBST and incubated with Alexa Fluor 488-conjugated secondary antibody. After washing with PBS, cells were mounted using VECTASHIELD Antifade Mounting Medium (Vector Laboratories, H-1000) or Prolong Gold Antifade Mountant with DAPI (Thermo Fisher, P36931). 2-dimensional images were acquired using a Leica SP8 confocal microscope. HA (or GFP) and LipidTOX Deep Red Neutral Lipid Stain were excited at 488 and 637 nm, and emission signals were collected at 495–535 and 645–700 nm, respectively. Bodipy 558/568 C_12_ was excited at 558nm and emission collected 565-600 nm.

### Western Blotting

Following treatments, cells were washed twice with ice cold D-PBS and proteins harvested using RIPA buffer supplemented with complete protease and PhosStop inhibitors (Sigma). The lysates were cleared by centrifugation at 13,000 rpm for 15 min at 4 °C, and protein concentration determined by a BioRad DC protein assay. Typically, 20-30 g of protein lysates were denatured in NuPAGE 4× LDS sample buffer and resolved on NuPage 4-12 % Bis-Tris gels (Invitrogen) and the proteins transferred by iBlot (Invitrogen) onto nitrocellulose membranes. The membranes were blocked with 5 % nonfat dry milk or 5 % BSA (Sigma) for 1 h at room temperature and incubated with the antibodies described below. Following a 16 h incubation at 4 °C, all membranes were washed five times in Tris-buffered saline-0.1% Tween-20 prior to incubation with horseradish peroxidase (HRP)-conjugated anti-rabbit immunoglobulin G (IgG), HRP-conjugated anti-mouse IgG (Cell Signaling Technologies). The bands were visualized using Immobilon Western Chemiluminescent HRP Substrate (Millipore). All images were acquired on the ImageQuant LAS 4000 (GE Healthcare). For mouse livers, protein was extracted from using T-PER Tissue Protein Extraction Reagent (Thermo Scientific #78510) containing freshly added 1x protease inhibitor (ApexBio # K10070) and 1x phosphatase inhibitor (Fisher Scientific #501905547). Nuclear protein was enriched using Abcam Nuclear Extraction Kit (#ab113474-100test) from snap frozen mouse liver tissue. The protein concentrations of all samples were measured using the Pierce BCA Protein Assay Kit (Life Technologies #23225). Samples were treated with 6x Laemmli SDS Sample buffer R (Boston BioProducts #BP-111R-25ml) and heated at 95 °C for 10 min. About 15 mg of protein per sample were separated using the BioRad 4∼20% gradient Tris-Glycine SDS mini gel system and analyzed using the antibodies indicated below.

### CIDEB structural modelling

AlphaFold predictions were carried out with AlphaFold2 ^39^ using the protocol developed for oligomers ^40^ on the Google <u>ColabFold v1.5.5</u> server. The quality of the prediction is described by the two main AlphaFold descriptors: the predicted local distance difference test (pLDDT) and predicted aligned error (PAE). The calculations were run for structures consisting of up to 12 monomeric chains. Only tetramers folded in the stable structure, reaching pLDDT 80-95 (confident to very high reliability). PAE descriptor indicates that all four protomers interact tightly in the tetramer (**Fig. S1**). Predictions of other oligomers led to structures of very low pLDDT (<40) and were discarded. The predictions of a tetramer were also run for human CIDEA and CIDEB and the most remote available sequence of hagfish CIDEB (UniProt entry A0A8C4N4T1) leading to very similar structures with root mean square errors of superposition of the b-sheet ranging from 1.9 to 2.4 to Å and comparable quality descriptors.

### Animal models

All mice were housed and handled in strict accordance with the guidelines of the Institutional Animal Care and Use Committee (IACUC) at UT Southwestern. *Cideb^−/-^* and *Cideb^fl/fl^* mice, maintained on the C57BL/6 background, were generated by the CRI Mouse Genome Engineering Core. For some experiments, *Cideb^fl/fl^* mice were crossed with *Alb-Cre* mice on the C57BL/6 background to achieve liver-specific embryonic deletion of *Cideb*. In general, experiments were initiated when mice were 6-8 weeks old. In most experiments, a high dose of AAV-TBG-Cre or AAV-TBG-GFP (5E10 GC/mouse) was administered to ensure efficient delivery to nearly all hepatocytes. Mouse sex is specified in the text or legends.

### Primary hepatocyte isolation and culture

Primary hepatocytes were isolated from mouse livers using a two-step collagenase perfusion method. Briefly, mice were anesthetized with isoflurane, and the abdominal area was sterilized with 70% ethanol. A midline incision was made to expose the portal vein, and a catheter was inserted into the inferior vena cava (IVC) and secured by threading a needle through the catheter’s butterfly wing. Perfusion was initiated with Liver Perfusion Medium (Life Technologies, #17701038) at a flow rate of 3 mL/min, using approximately 30–50 mL of medium. The portal vein was immediately severed to allow for adequate outflow. Following perfusion with Liver Perfusion Medium, the liver was perfused with Liver Digestion Medium (Life Technologies, #17703034) at the same flow rate, using an additional 30–50 mL. After digestion, the liver was carefully removed and placed in 10 mL of Liver Digestion Medium in a 10 cm culture dish. The liver capsule was gently peeled away, and the tissue was gently agitated to release hepatocytes. The cell suspension was then filtered through a 70 µm nylon mesh into a 50 mL tube prefilled with 20 mL of Hepatocyte Washing Medium (Life Technologies #17704024) on ice to halt enzymatic activity. Cells were pelleted by centrifugation at 50 g for 5 minutes at 4°C, and the supernatant was discarded. The cell pellet was resuspended in 20 mL of Washing Medium, and the wash step was repeated twice to ensure cell purity. Cell viability and yield were assessed by Trypan blue exclusion. Cells were then centrifuged once more at 50 g for 5 minutes and resuspended in the desired volume of washing medium or buffer for further experiments.

### FFA treatment assay

Human liver cancer cell lines (HepG2 and Huh7), a mouse immortalized liver cell line (H2.35), and primary hepatocytes (*Cideb^fl/fl^* and *Alb-Cre*; *Cideb^fl/fl^*) were cultured and treated with saturated and unsaturated fatty acids. Cells were exposed to fatty acids at a final concentration of 100 µM for 16 hours. Saturated fatty acids included palmitic acid (16:0) (Sigma Aldrich #43051), stearic acid (18:0) (Sigma Aldrich #85679), and arachidic acid (20:0) (Sigma Aldrich #A3631). Unsaturated fatty acids included oleic acid (18:1) (Sigma Aldrich #O1257), linolenic acid (18:3) (Sigma Aldrich #L1376), stearidonic acid (18:4) (Sigma Aldrich #SMB00291), mead acid (20:3) (Sigma Aldrich #43059), arachidonic acid (20:4) (Sigma Aldrich #23401), and eicosapentaenoic acid (20:5) (Sigma Aldrich #E3635). All fatty acids (50 mM) were dissolved in ethanol, then diluted to 5 mM with warm 10% fatty acid-free BSA (Sigma Aldrich #A8806) prior to application to cells to ensure optimal cellular uptake. After incubation, cells were harvested for further analysis as described in subsequent sections.

### Dietary models of MASLD and ALD

WD consisted of high sugar, high fat, and high cholesterol solid food (Teklad Diets #TD. 120528) combined with high-sugar water containing 23.1 g/L d-fructose (Sigma-Aldrich #G8270) and 18.9 g/L d-glucose (Sigma-Aldrich #F0127). CDA-HFD consisted of a methionine and choline-deficient high-fat diet (Research Diets #A06071302). The NIAAA alcohol feeding protocol is comprised of ethanol liquid diet (1-5%, vol/vol), and 1000 mL of ethanol liquid diet add to a blender 133 g of dry mix (Bio-serv, Product F1258SP) and different amounts of maltose dextrin and a different volume of water to the diet on the basis of the percentage of ethanol required ^23^ with weekly oral gavage of 5 μg/kg ethanol.

### Plasma and liver metabolic assays

At the study endpoints, blood was collected using heparinized tubes (Fisherbrand #22-362-566) from the inferior vena cava immediately after sacrificing the mouse. Blood was then transferred into 1.5 mL tubes and centrifuged at 2000 g for 15 minutes at 4 °C. The supernatant (plasma) was analyzed for AST, ALT, cholesterol, and triglyceride using specific reagent kits (VITROS 8433815, 1655281, 1669829, 1336544) on a fully automated OCD Vitros 350 dry chemistry analyzer. All analyses followed the protocols provided by the reagent kit manufacturer (Ortho Clinical Diagnostics, Raritan, NJ) at the UT Southwestern Metabolic Phenotyping Core. BHB was measured in the plasma following manufacturer’s instructions (Cayman #700190). Dilutions were 1:3 for mouse plasma. To measure cholesterol and triglycerides in liver tissues ^41^, 100–150 mg of liver tissue is homogenized in 4 ml of a 2:1 chloroform:methanol solution. After adding 800 µl of saline, the mixture was vortexed, centrifuged, and the organic layer was collected and transferred into a 5 ml volumetric flask, topped off with chloroform to the exact 5 ml mark, and left overnight. The next day, samples and standards were prepared in a 96-well plate with a Triton-chloroform mix, vortexed, and dried. On the final day, cholesterol or triglyceride reagents were added, the plate was incubated at 37°C for 15 minutes, and absorbance was measured to quantify the lipid content. Values were normalized to sample weight.

### Histology

Tissue samples were fixed overnight at 4°C in 4% paraformaldehyde. Fixed tissues were embedded in paraffin, sectioned and H&E staining by the UTSW Histopathology Core or the SCCC Tissue Management Service. Fibrosis detection was performed by staining with Sirius Red (IHC World #IW-3012) and quantified by ImageJ. Images were taken by a Hamamatsu Nanozoomer 2.0HT in the UTSW Whole Brain Microscopy Facility.

### AAV production and purification

AAV-Pro 293T cells (Takara #632273) were plated one day before transfection at 50% confluence, allowing the cells to reach 80-90% confluence the next day. For transfection of one 15 cm dish, the following plasmids were mixed in one tube with 1 mL Opti-MEM medium: 10 μg MOSAICS-V8 or -V10 vector, 10 μg pAAV2/8 (Addgene #112864), and 20 μg pAdDeltaF6 (Addgene #112867). In another tube, 160 μl PEI solution (1 mg/mL in water, pH 7.0, ChemCruz #sc-360988) was mixed with 1 mL Opti-MEM medium. The solutions from both tubes were then combined and incubated for 10 minutes before being added to the cell culture. Forty-eight hours after transfection, the cells were scraped off the dish and collected by centrifugation at 500 g for 10 minutes. The supernatant was disinfected and discarded, and the cell pellets were lysed in 1.5 mL lysis buffer per 15 cm dish (PBS supplemented with NaCl to a final concentration of 200 mM and CHAPS to a final concentration of 0.5% (w/v)). The cell suspension was placed on ice for 10 minutes with intermittent vortexing, and then centrifuged at 20,000 g for 10 minutes at 4°C. The supernatant containing the AAV was collected. To set up the gravity column for AAV purification, 0.5 mL of AAV8-binding slurry beads (ThermoFisher #A30789) was loaded into an empty column (Bio-Rad #731-1550). After the beads were tightly packed at the bottom, they were washed with 5 mL wash buffer (PBS supplemented with NaCl to a final concentration of 500 mM). The supernatant containing AAV was then loaded onto the column. After all the supernatant flowed through, the beads were washed twice with 10 mL wash buffer. The AAV was then eluted with 3 mL elution buffer (100 mM glycine, 500 mM NaCl in water, pH 2.5) and the eluate was immediately neutralized with 0.12 mL 1 M Tris-HCl (pH 7.5-8.0). The AAV was concentrated by centrifugation at 2000 g for 3-5 minutes at 4°C using a 100k Amicon Ultra Centrifugal Filter Unit (Millipore #UFC810024). After centrifugation, the volume of AAV was adjusted to equal or less than 0.5 mL. The concentrated AAV was diluted with 4-5 mL AAV dialysis buffer (PBS supplemented with NaCl to a final concentration of 212 mM and 5% sorbitol (w/v)) and centrifuged at 2000 g for 3-5 minutes at 4°C. The dilution and centrifugation processes were repeated three times. The final concentrated AAV was transferred into a 1.5 mL tube and centrifuged at 20,000 g for 5 minutes to remove debris. The supernatant was aliquoted, flash frozen using liquid nitrogen, and stored at -80°C. The AAV titer was determined using the AAVpro Titration Kit (for Real Time PCR) Ver.2 (Takara Bio #6233).

### MOSAICS screening and Cas9 mediated gene deletion in mice

SgRNAs (see Table sgRNA primers) were cloned into the MOSAICS-V8 or -V10 vectors, and the corresponding AAV8 was produced. For single gene deletion, AAV8 at a concentration of 2E11 VG/mL was diluted to a final volume of 100 μl with saline. This solution was retro-orbitally injected into *Rosa-rtTA; TetO-Cas9* double homozygous mice (iCas9) 3-5 days after initiating doxycycline (dox) water (1 mg/mL dox) at 6.5 weeks of age. For the AAV libraries used in the LD screens, AAV8 at a concentration of 5E11 VG/mL was injected. For liver-wide single or double gene deletion, we used AAV8 at a concentration of 1E12 VG/mL. One week post-injection, the mice were fed NC, WD, or CDA-HFD. The reads from the sequencing of amplicon libraries were processed using cutadapt (version 1.9.1) to remove excessive adaptor sequences, retaining only the sgRNA sequences. The 5’ sequences were trimmed using the options ‘-O 32 –discard-untrimmed -g CTTTATATATCTTGTGGAAAGGACGAAACACCG’, and the 3’ sequences were trimmed using the options ‘-O 12 -a GTTTTAGAGCTAGAAATAGCÀ. The abundance of each sgRNA was calculated with the count function in MAGeCK (version 0.5.6) using default options. The trimmed fastq files were assigned to NC, WD, and CDA-HFD groups and uploaded, along with library files containing sgRNA sequences and targeted gene names, to a server preloaded with MAGeCK. The enrichment of each sgRNA was calculated using the test function in MAGeCK.

### Genomic DNA extraction, sgRNA amplification, and amplicon library construction

To extract genomic DNA containing the integrated sgRNA, the entire liver was minced into approximately 1 mm³ pieces using a blade and weighed. Minced liver was transferred into a glass Wheaton Dounce Tissue Grinder in two volumes (w/v) of homogenizing buffer (100 mM NaCl, 25 mM EDTA, 0.5% SDS, 10 mM Tris-HCl, pH 8) and stroked 50 times or until no bulk tissue remained. After homogenizing, 500 μl liver lysate was transferred to a 15 mL tube for genomic DNA extraction using the Blood & Cell Culture DNA Midi Kit (Qiagen #13343) according to the manufacturer’s protocol. The remaining lysates were frozen at -80°C as backup samples. For sgRNA amplification and amplicon library construction, 5 μg of genomic DNA, 5 μl of general forward primer mix (5 mM), 5 μl of barcode-specific reverse primer (5 mM), 1 μl of Q5 DNA polymerase, 10 μl of Q5 buffer, 10 μl of High GC buffer, 1 μl of dNTP (10 mM), and water were mixed for a 50 μl PCR reaction. Two reactions were made for each genomic DNA sample. The PCR cycle was: 95°C for 3 min, followed by (95°C for 30 s, 60°C for 30 s, and 72°C for 20 s) repeated n times, and a final extension at 72°C for 2 min. The cycle number was pre-optimized using the same PCR reactions with a smaller volume, choosing the cycle number that gave a weak but sharp band on a DNA gel. For the final PCR reaction, 25 cycles were used. After PCR, 50 μl was resolved on a DNA gel. The 250 bp band corresponding to the amplicon was cut and purified using the QIAquick Gel Extraction Kit (Qiagen #28704). DNA concentration was determined using the Qubit kit (Invitrogen #Q32853), and high-throughput sequencing was performed using an Illumina NextSeq500 system at the CRI at UT Southwestern Sequencing Facility ^16,42^.

### RNA-seq analysis

Total RNA was purified using Trizol and the Qiagen RNeasy Mini Kit (Qiagen #74104). An RNase-free DNase Set (Qiagen #79254) was employed for on-column treatment to remove any trace amounts of genomic DNA. Total RNA samples that passed quality control—having an RNA integrity number above 7, as determined by Agilent’s RNA ScreenTape Analysis (Agilent #50675576)—were used for RNA-seq library preparation. This was performed with the NuGen Ovation Mouse RNA-Seq Systems 1–16 (Nugen #0348-32) and sequenced with 75 bp single-end flow cells using an Illumina NextSeq500 system at the Children’s Research Institute sequencing core. RNA-seq data were analyzed using the nf-core/rnaseq pipeline. Sequencing adaptors were removed with Trim Galore (v0.6.7) [EMBnet.journal: https://doi.org/10.14806/ej.17.1.200]. Sequencing reads were mapped to the mouse genome assembly GRCm39 (mm39) using the STAR aligner (v2.7.9a). Gene expression was quantified with Salmon (v1.10.1). Differential gene expression analysis was conducted using the R package DESeq2 within the nf-core/differential abundance pipeline. Genes were considered differentially expressed if their log2 fold changes were greater than 1 and their adjusted p-values were less than 0.05. Z-scores of variance-stabilizing-transform normalized counts were used to plot. For WD and CDA-HFD RNA-seq data, alignment, quantification, and differential expression analysis were performed using the QBRC_BulkRnaSeqDE pipeline (https://github.com/QBRC/QBRC_BulkRnaSeqDE). Briefly, the alignment of reads to the mouse reference genome GRCm38 (mm10) was done using STAR aligner (v2.7.2b).31 Gene count quantification was done using FeatureCounts (v1.6.4).32 Differential gene expression analysis was performed using the R package DEseq2 (v1.26).33 Cutoff values of absolute fold change greater than 2 and FDR<0.05 were used to select for differentially expressed genes between sample group comparisons. Finally, GSEA was carried out with the R package fgsea (v1.14.0) using the ‘KEGG’ and ‘Hallmark’ libraries from MsigDB.

### RNA extraction and real time qPCR

Total RNA was isolated from liver tissue using TRIzol reagent (Invitrogen #15596018), followed by purification with the RNeasy Mini kit (Qiagen #74104). For real-time quantitative PCR, cDNA synthesis was performed from 1 μg of total RNA using the iScript Reverse Transcription Supermix (BioRad #1708840) in a 20 μL reaction. Each resulting cDNA sample was then diluted to a final volume of 200 μL. For PCR, 2 μL of this diluted cDNA was combined with specific primers and iTaq Universal SYBR Green Supermix (BioRad #1725121) in a 10 μL reaction.

### Poloxamer-407 assay

Mice were fasted overnight before initiation of the VLDL secretion assay. After baseline blood collection (∼50 μl) from the retro-orbital sinus, mice were intraperitoneally injected with poloxamer-407 (P-407) (10% (w/v)) in saline (Sigma-Aldrich, #16758) at a dose of 1.0 g/kg of body weight. Blood samples (approximately 50 μL) were collected at 0, 90, 180 min post-injection. Plasma was separated by centrifugation and triglycerides were measured as described above.

### FFA analysis

Approximately 20 mg of frozen liver tissue was homogenized in a 2:1:1 mixture of chloroform, PBS, and methanol. After homogenization, samples were centrifuged in screw-cap glass vials at 1,000 g at 4°C for 10 minutes. The chloroform layer was transferred to a new vial and dried under nitrogen. Dried samples were saponified with 1 M KOH in 80% methanol at 56°C for 1 hour, neutralized with HCl, and re-dried under nitrogen. Samples were then re-suspended in a 65:35 isopropanol/methanol mixture for LC-MS/MS analysis. Fatty acids were resolved on a Thermo Scientific Vanquish Flex UHPLC system with an Accucore C18 column (2.1×150 mm), using a binary solvent system (A: 50:50 water:acetonitrile; B: 88:10:2 isopropanol:acetonitrile:water with 5 mM ammonium formate and 0.1% formic acid). A programmed gradient flow (0.17 mL/min) separated fatty acids, with the column at 60°C. Mass spectrometry was performed on a Thermo Fusion Lumos Tribrid Orbitrap, using MSn acquisition for structural identification of lipid species ^27^. Data were analyzed with TraceFinder software (Thermo Fisher Scientific, V5.0) using a validated library of fatty acid spectra.

### Isotope tracing to assess DNL and FAO

Mice received an intraperitoneal injection of 99.9% D_2_O (Sigma-Aldrich, 151882) in 0.9% NaCl at 30 µL/g body weight. After 5 hours, mice were sacrificed and ∼20 mg liver samples were homogenized in a 2:1:1 mixture of chloroform, PBS, and methanol, then centrifuged at 1,000 g at 4°C for 10 minutes. The chloroform layer was dried under nitrogen, saponified with 1 M KOH in 80% methanol at 56°C for 1 hour, neutralized with HCl, re-dried, and re-suspended in a 65:35 isopropanol/methanol mixture for analysis. Mass spectrometry was performed on a Thermo Fusion Lumos Tribrid Orbitrap. Extracted ion chromatograms for labeled and non-labeled species were generated with 1-5 ppm mass accuracy to distinguish ^13^C and ^2^H isotopes. Data analysis used TraceFinder software (V5.0), with isotope correction via a customized R script based on the AccuCor algorithm. For FAO, isotope tracing was conducted with [U-^13^C]linolenic acid or [U-^13^C]stearic acid (Cambridge Isotope Laboratories, CLM-6990-PK and CLM-8386-1) dissolved in ethanol at 50 mM, diluted to 5 mM with warm 10% fatty acid-free BSA. Mice were infused at 1 mg/min/kg for 2 hours. Approximately 20 mg of frozen liver tissue was homogenized in 80:20 acetonitrile:water, subjected to freeze-thaw cycles, and centrifuged at 16,000 g for 15 minutes at 4°C. The supernatant was analyzed on a Thermo Scientific QExactive HF-X hybrid quadrupole orbitrap HRMS ^28^. Data analysis was performed with TraceFinder software (V5.0), with isotope abundances corrected via a customized R script based on the AccuCor algorithm ^43^.

### The fatty acid oxidation assay

The fatty acid oxidation assay was conducted using [1-^14^C]-stearic acid isotope tracing in hepatocytes cultured to 90% confluence in 6-well plates. Cells were incubated in DMEM containing 10% FBS, 50 μM ^12^C-stearic acid (Sigma Aldrich #85679), 2 μCi [1-^14^C]-stearic acid (American Radiolabeled Chemicals, ARC0258), 0.28% BSA (Goldbio, A-421-10), and 1 mM L-carnitine (Millipore Sigma, C0283). Prior to incubation, [1-^14^C]-stearic acid was dried at 37°C overnight and conjugated with BSA and 12C-stearic acid. Hepatocytes were exposed to this mixture for 6 hours. Released ^14^CO_2_ was captured on filter paper pre-soaked with 30 μl of 1N NaOH for 2 hours. The filter paper was then transferred to a scintillation vial (VWR, 66022-055) for analysis. The CPM value was measured for 1 minute using a scintillation counter (Beckman Coulter LS 6500 Liquid Scintillation Counter). For experiments involving linolenic acid (American Radiolabeled Chemicals, ARC0379), the same protocol was followed.

### Statistical analysis

The data in most panels reflect multiple experiments performed on different days using mice derived from different litters. In all experiments, mice were not excluded from analysis after the experiment was initiated unless the mice died. Unless otherwise stated in the methods or figure legends, 2-tailed Student’s *t* tests (2-sample equal variance) were used to test the significance of differences between 2 groups. For time-course–expression experiments, 1-way ANOVA was performed and the significance shown was compared with the initial time point. For experiments involving 2 groups and different time points, 2-way ANOVAs were used, and the significance was compared with the means of each group at different time points. Variation is indicated using SEM and presented as mean ± SEM. Statistical significance was defined as *P* < 0.05.

### Liver cancer modelling

For the mutagen induced HCC model, (50 μg/g of DEN (Sigma #N0756) was given at p14 to both male and female mice. At 28 days of age, these mice were injected with either AAV-TBG-GFP or AAV-TBG-Cre and initiated on WD. After 8 months, livers were harvested.

---

**Fig. S1.**
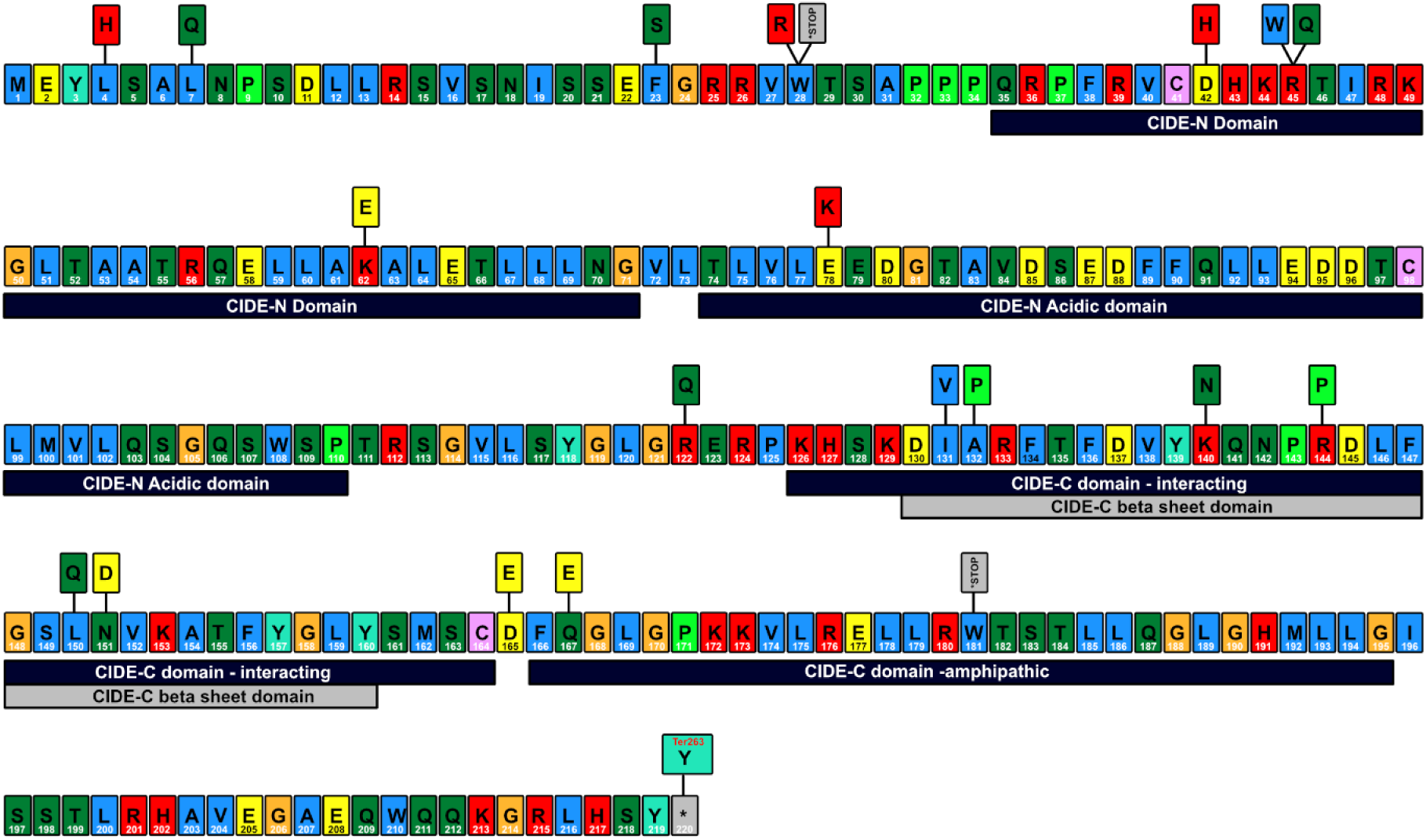
Somatic *CIDEB* mutations in patients with chronic liver disease. Distribution of somatic mutations in *CIDEB*. Amino acids with observed mutations in chronic liver disease shown above the WT protein sequence

**Fig. S2.**
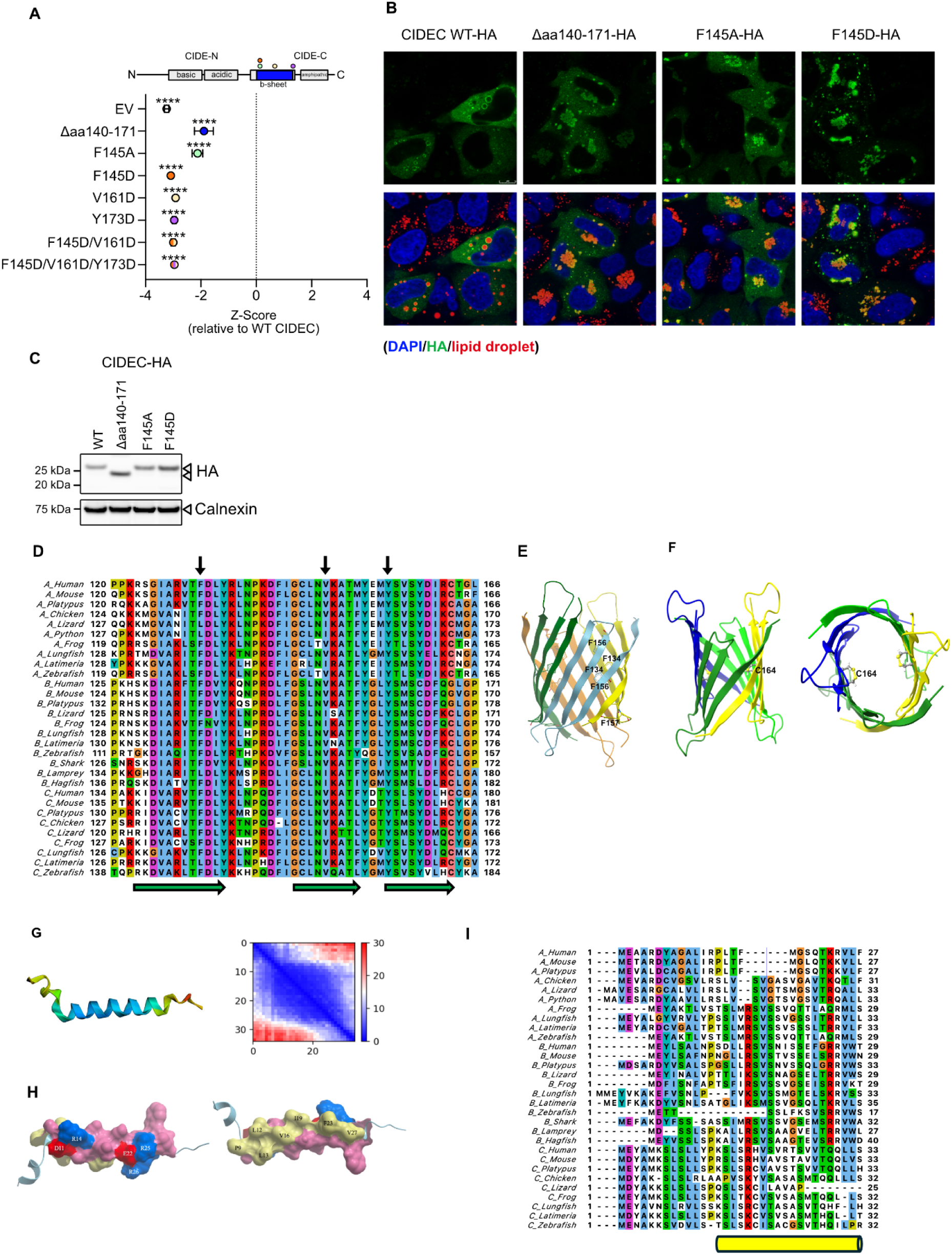
Disruption of the predicted C-terminal β-sheet and N-terminus helix on CIDEC lipid droplet enlargement activity. **A.** Quantification of LD volume in oleate-loaded HeLa cells transfected with CIDEC β-sheet deletion and point mutants. Enlargement activity is represented as a Z-score where 0 is WT CIDEC. **B.** Representative images of LDs (red) and HA (green) in oleate-loaded HeLa cells transfected with CIDEC β-sheet deletion and point mutants. Nuclei were stained with DAPI (blue). **C.** Western blot analysis in oleate-loaded HeLa cells transfected with CIDEC β-sheet deletion and point mutants. **D.** Sequence alignment of the β-sheet region of CIDE proteins from representative vertebrates. β-sheet strands from AlphaFold predicted tetramers are indicated by green arrows. The position of the hydrophobic residues within the β-sheet (indicated by arrows), the mutation of which to acidic D impaired LD growth. A, B, and C stand for CIDEA, CIDEB and CIDEC, respectively. **E.** Cluster of aromatic residues inside the CIDEB putative channel (hollow β-barrel). **F.** C164 in two antiparallel protomers of the CIDEB channel can form disulfide bonds. Side view (left panel) and along the axis (right panel) of the channel. **G.** AlphaFold prediction of the N-terminal segment of CIDEB aa 1-34 with quality parameters coloured as in Fig. 2A. **H.** Surface representation of the helix, predicted with confidence, viewed from two sides and coloured according to residue type (yellow: hydrophobic, red: negative, blue: positive, violet: others). **I.** Sequence alignment of the N-terminal region of CIDE proteins from representative vertebrates. The α-helix predicted by AlphaFold in human CIDEB is indicated by a yellow cylinder. The insertion between positions 16-28 (blue line), unique to B_Hagfish, is not shown. All data are presented as mean ± SEM ( ****p < 0.0001).

**Fig. S3.**
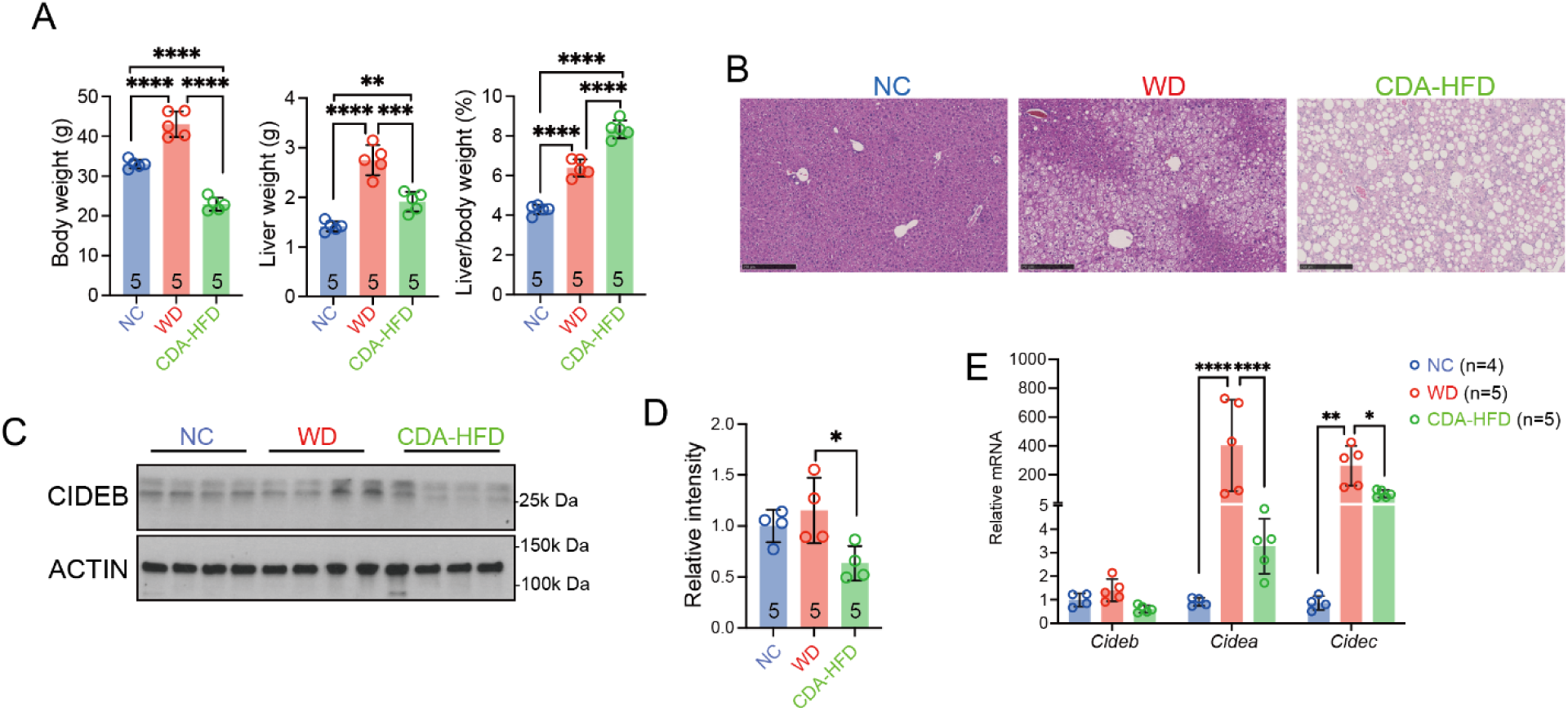
*CIDEB* expression in different MAFLD models. **A.** Body weight, liver weight, and liver/body weight ratio in C57BL/6 mice fed NC, WD, or CDA-HFD for 12 weeks. **B.** Representative H&E staining of livers treated with NC, WD, or CDA-HFD for 12 weeks. **C.** Western blot analysis to assess CIDEB levels in the liver. **D.** Quantification of CIDEB protein levels. **E.** qPCR for *Cideb*, *Cidea*, and *Cidec* mRNA in livers fed with NC, WD, or CDA-HFD. All data are presented as mean ± SEM (*p < 0.05; **p < 0.01; ***p < 0.001; ****p < 0.0001).

**Fig. S4.**
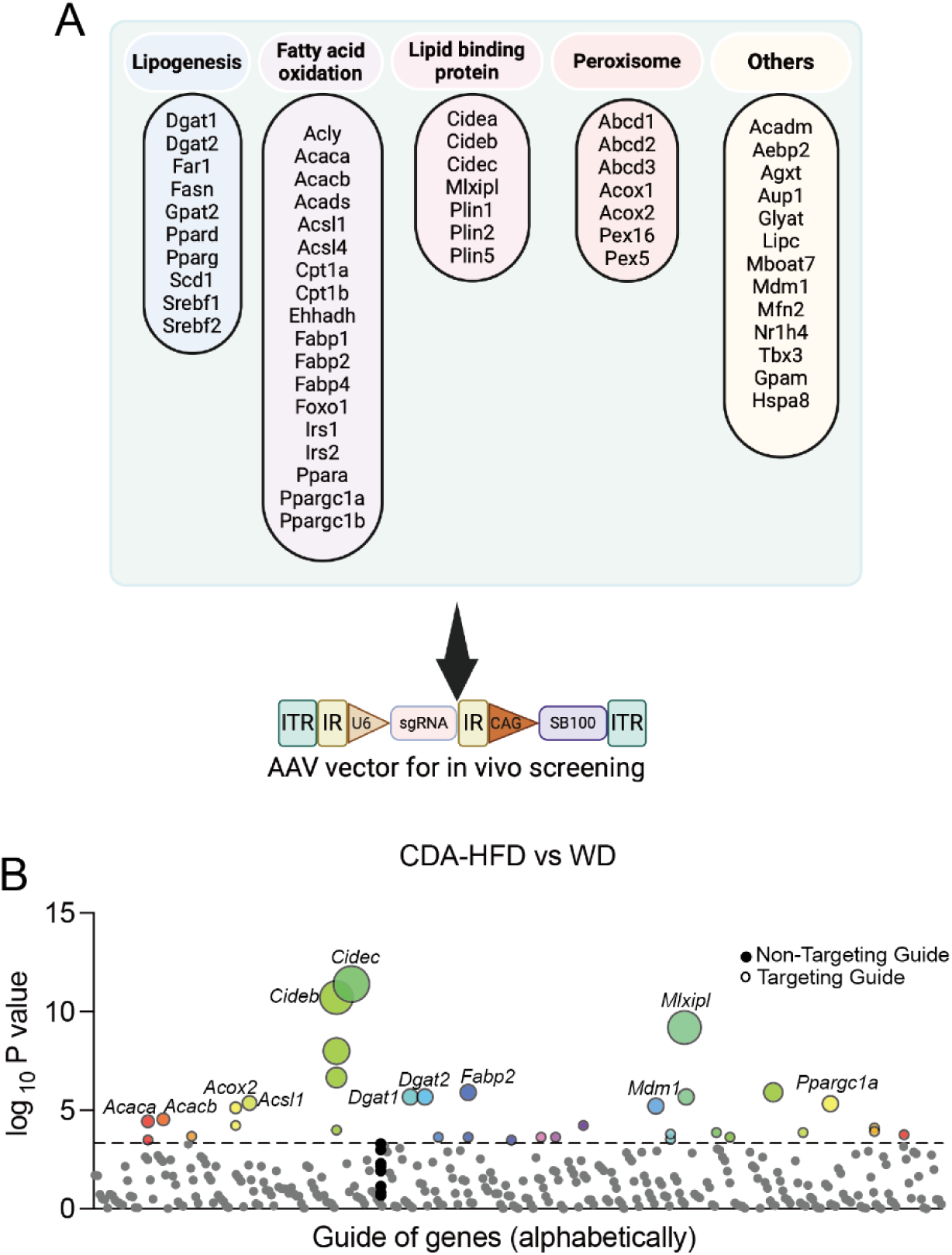
In vivo CRISPR screening of genes involved in fatty liver disease. **A.** Genes targeted in the CRISPR screen. **B.** sgRNAs enriched in CDA-HFD or WD but not in NC-fed livers. Each circle represents one sgRNA. Different sgRNAs targeting the same genes were aligned vertically. Circle sizes correlate to -log_2_(p). Control sgRNAs are filled black circles.

**Fig. S5.**
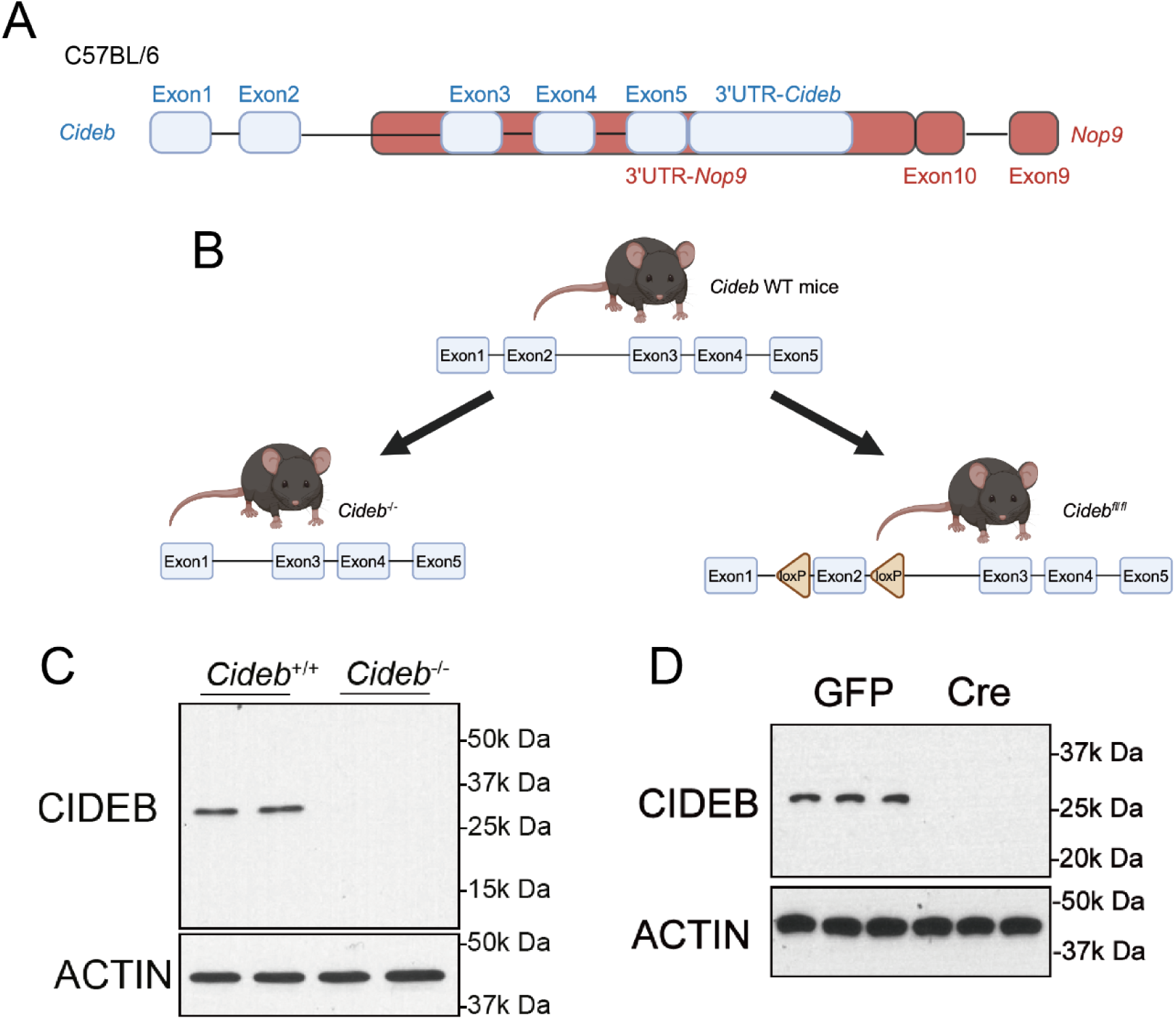
Generation and validation of *Cideb* KO mice. **A.** Schematic representation of the mouse *Cideb* gene and overlap with the *Nop9* gene. **B.** Strategy to generate whole-body *Cideb* KO and floxed mice. **C.** CIDEB protein in WT and whole-body KO livers. **D.** CIDEB protein in livers from *Cideb*^fl/fl^ mice injected with AAV-TBG-GFP or AAV-TBG-Cre.

**Fig. S6.**
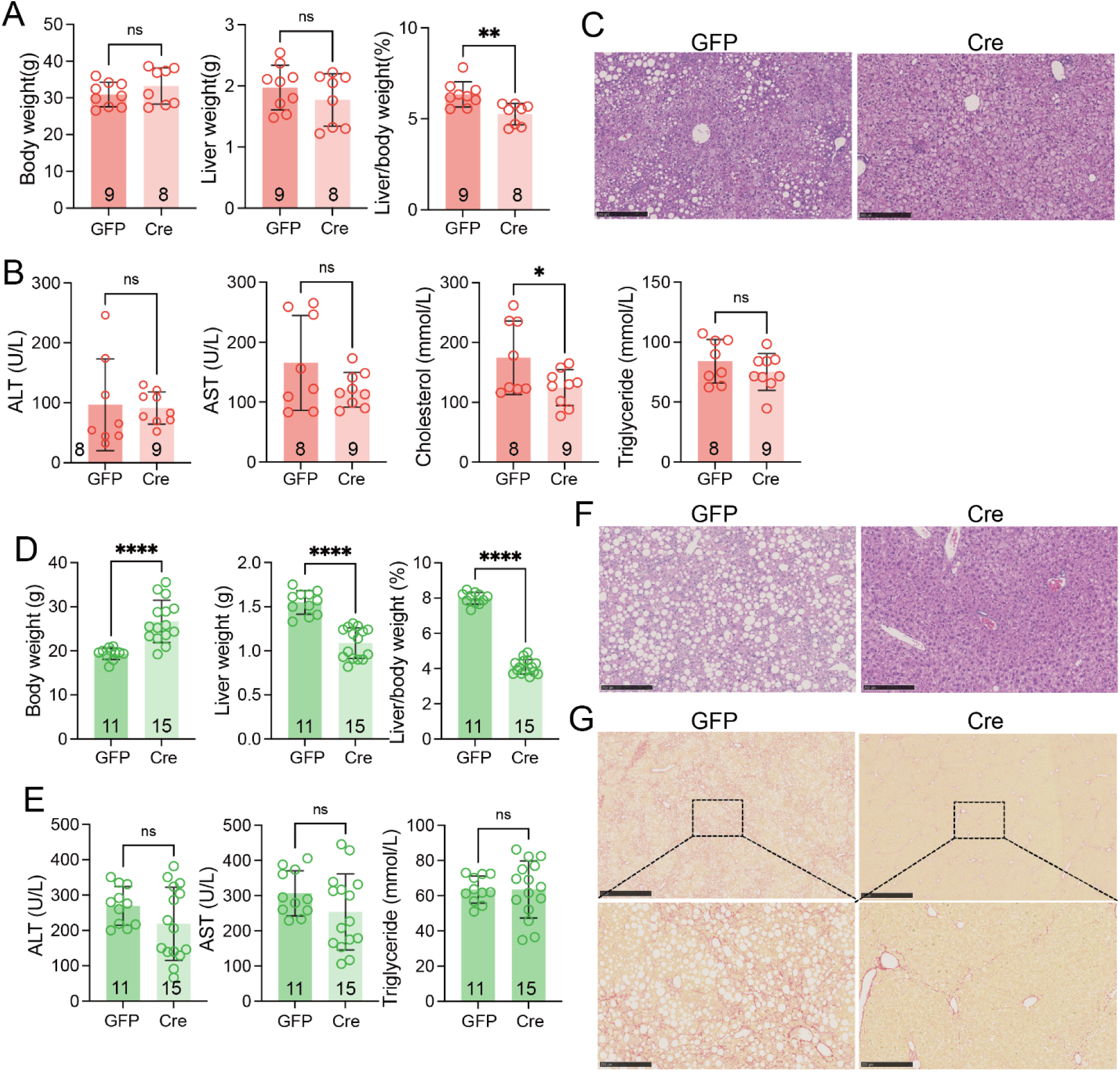
Impact of hepatocyte-specific *Cideb* deletion in female mice. **A.** Body weight, liver weight, and liver-to-body weight ratio in mice injected with AAV-TBG-GFP or AAV-TBG-Cre and fed with 24 weeks of WD. **B.** Plasma ALT, AST, cholesterol, and triglycerides in WD fed mice. **C.** Representative H&E staining of livers from **Fig.S6A**. **D.** Body weight, liver weight, and liver-to-body weight ratio in mice injected with AAV-TBG-GFP or AAV-TBG-Cre and fed with 12 weeks of CDA-HFD. **E.** Plasma ALT, AST and triglycerides in CDA-HFD fed mice. **F.** Representative H&E staining of livers from **Fig.S6D**. **G.** Representative Sirius red staining of livers from **Fig.S6D**. All data are presented as mean ± SEM (*p < 0.05; **p < 0.01; ****p < 0.0001).

**Fig. S7.**
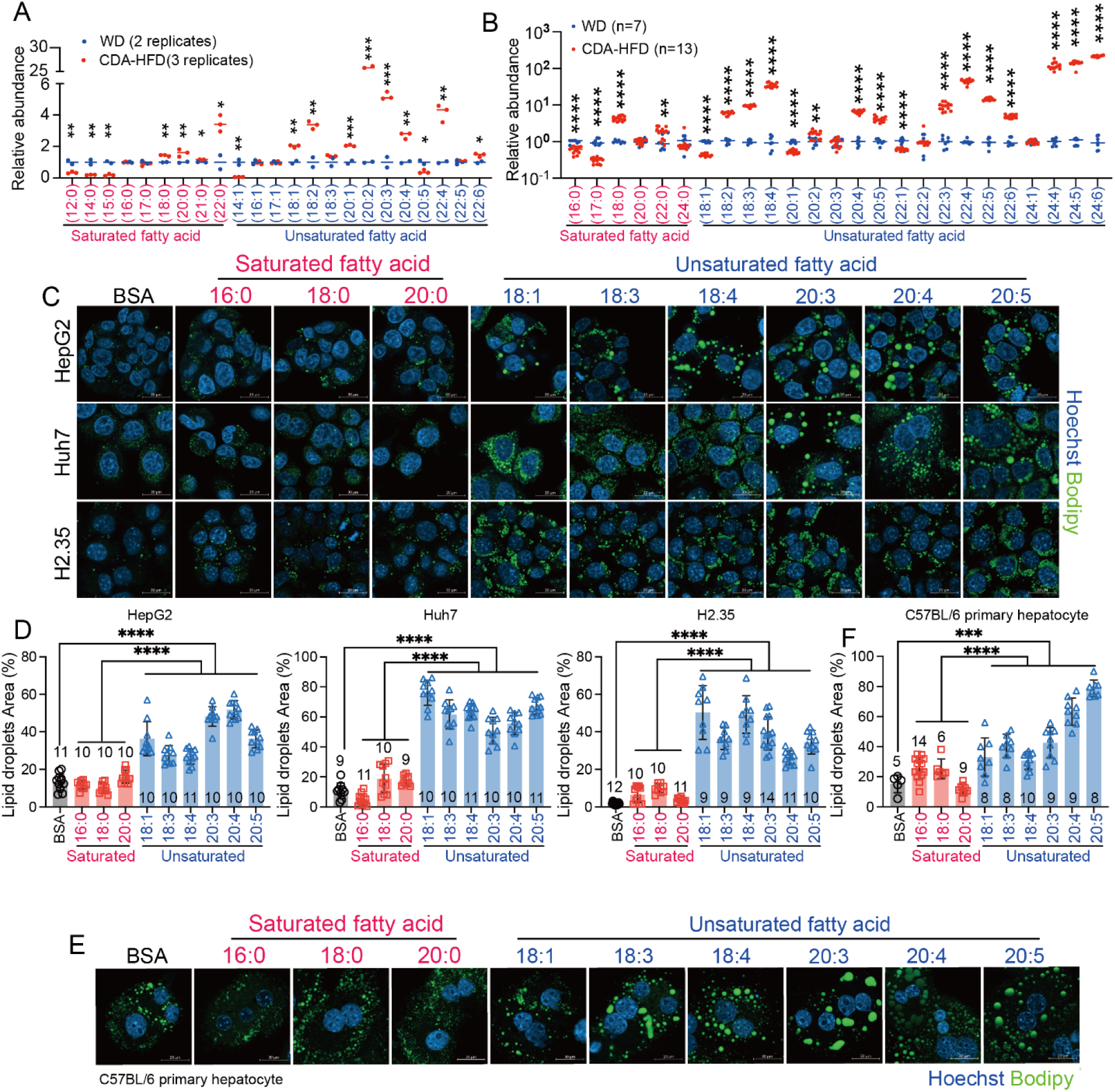
Unsaturated fatty acids preferentially accumulate within LDs in vitro and in vivo. **A.** Lipidomic analysis of the fatty acid content of WD and CDA-HFD. **B.** Lipidomic analysis of saturated and unsaturated fatty acids in WT livers exposed to WD or CDA-HFD for 12 weeks. **C.** Representative images of LD staining (green) of HepG2, Huh7, and H2.35 cells treated with 10% BSA, saturated (16:0, 18:0, 20:0) or unsaturated fatty acids (18:1, 18:3, 18:4, 20:3, 20:4, 20:5) at a concentration of 100 µM for 16 hours. LDs were stained with Bodipy (green) and nuclei were stained with Hoechst (blue) (scale bars = 20 µm). **D.** Quantification of the LD area (%) in the cells from C under fatty acid treatments. **E.** Representative images of LD staining (green) of C57BL/6 primary hepatocytes treated with various lipids, same as in **Fig.S7C**. **F.** Quantification of the LD area (%) from **Fig.S7E**. All data are presented as mean ± SEM (*p < 0.05; **p < 0.01; ***p < 0.001; ****p < 0.0001).

**Fig. S8.**
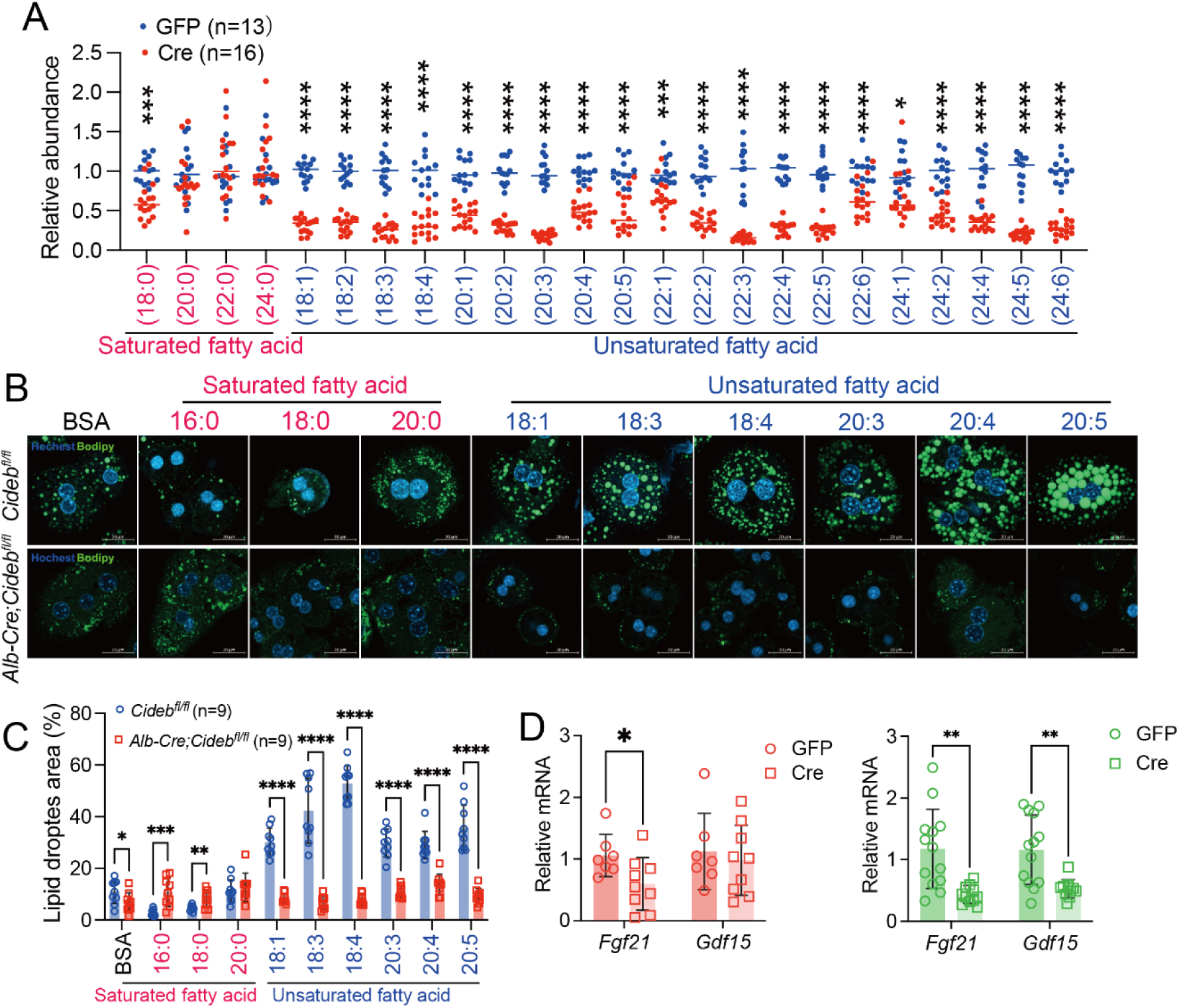
*Cideb* deletion reduces unsaturated fatty acids and LD growth in the liver. **A.** Relative abundance of fatty acids in *Cideb*^fl/fl^ livers injected with AAV-TBG-GFP or AAV-TBG-Cre and fed with 12 weeks of CDA-HFD. **B.** Representative LD staining (green) of primary hepatocytes from *Cideb*^fl/fl^ and *Alb-Cre; Cideb^fl/fl^* mice (scale bars = 20 µm). **C.** Quantification of LD area from **Fig.S8B**. **D.** qPCR of *Gdf15* and *Fgf21* from WT and whole body KO livers of CDA-HFD treated mice. All data are presented as mean ± SEM (*p < 0.05; **p < 0.01; ***p < 0.001; ****p < 0.0001).

**Fig. S9.**
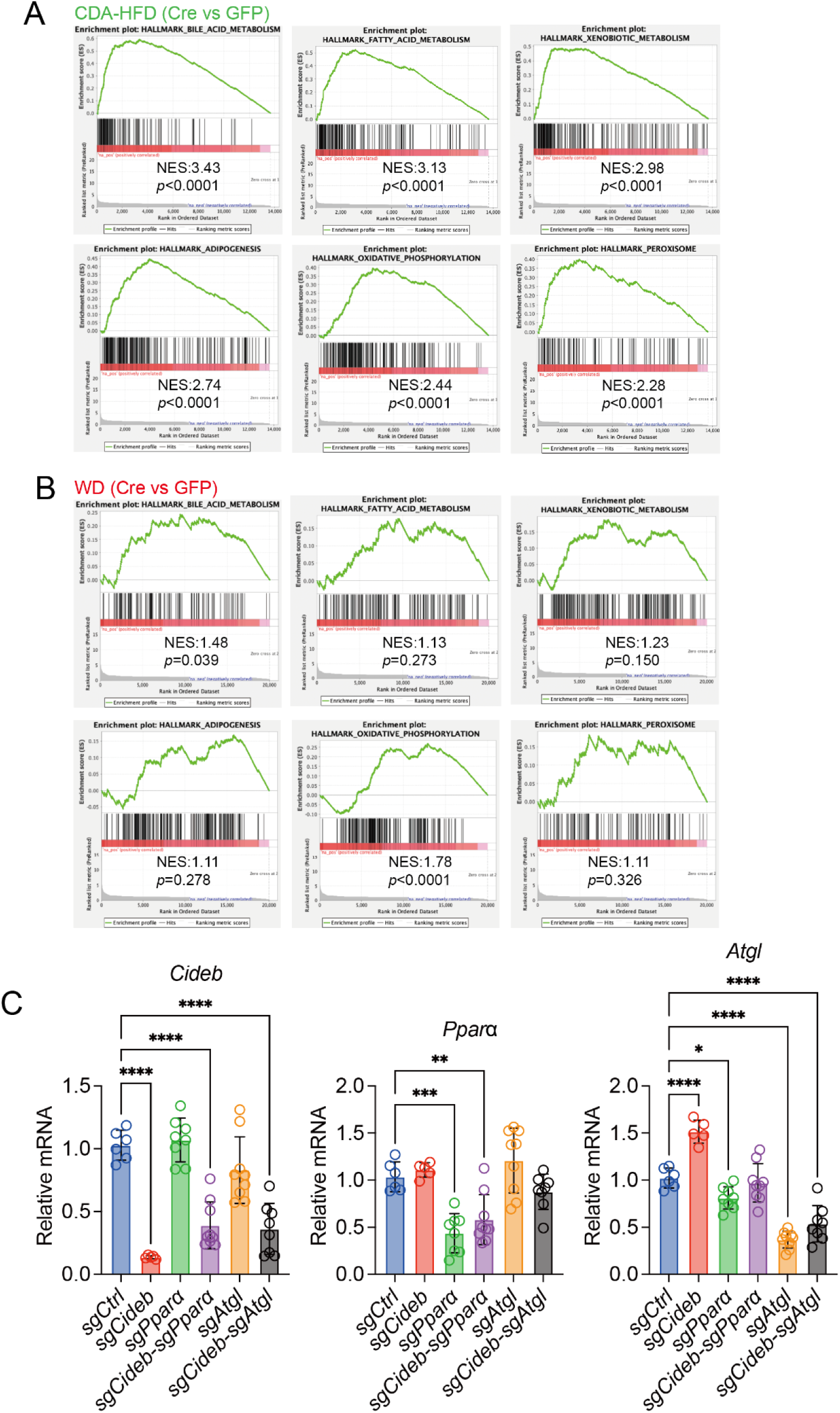
*Cideb* deletion alters liver gene expression in the context of MAFLD inducing diets. **A.** Gene Set Enrichment Analysis (GSEA) of RNA-seq data from 12 week CDA-HFD treated *Cideb^fl/fl^* livers injected with either AAV-TBG-GFP or AAV-TBG-Cre. **B.** GSEA of RNA-seq data from 24 week WD treated *Cideb^fl/fl^* livers injected with either AAV-TBG-GFP or AAV-TBG-Cre. **C.** qPCR of *Cideb, Pparα* and *Atgl* from iCas9 mice given high-dose AAV-sgRNAs (*sgCtrl*, *sgCideb*, *sgPparα*, *sgCideb*/*sgPparα, sgAtgl or sgCideb/sgAtgl*) treated with CDA-HFD for 3 weeks (n=6, 5, 8, 9, 9 and 8 mice). All data are presented as mean ± SEM (*p < 0.05; **p < 0.01; ***p < 0.001; ****p < 0.0001).

**Fig. S10.**
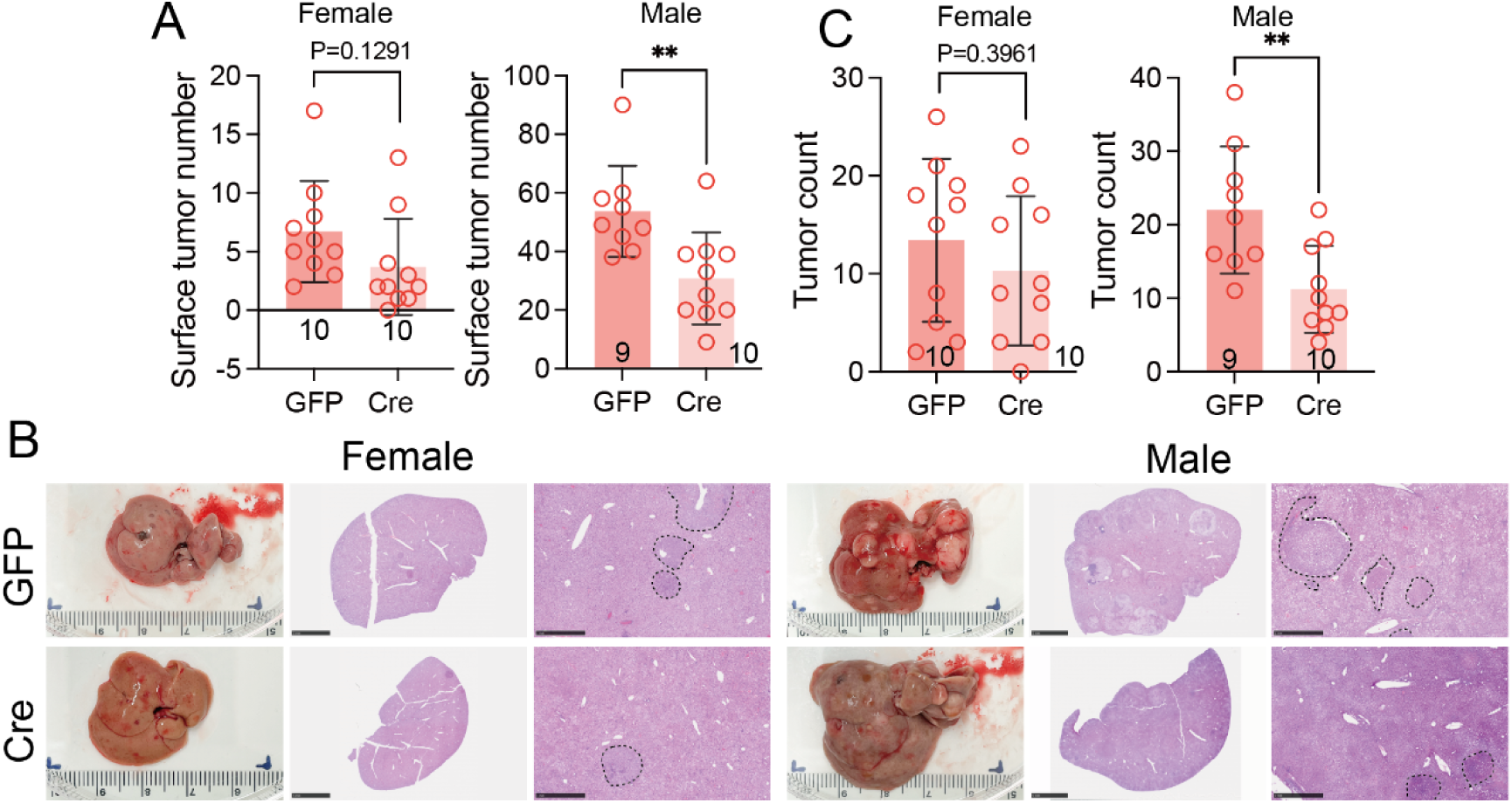
*Cideb* deletion reduces liver tumor development in male mice. **A.** Surface tumor numbers in DEN-treated livers. *Cideb* was deleted using AAV-TBG-Cre at P28. **B.** Representative picture and H&E staining of DEN treated livers from **Fig.S10A**. **C.** Quantification of tumor nodules. All data are presented as mean ± SEM. **p < 0.01.

